# Aging-associated metabolomic and lipidomic remodeling in mouse brain endothelial cell senescence *in vitro*

**DOI:** 10.1101/2025.08.14.670255

**Authors:** Ammar Tahir, Hari Baskar Balasubramanian, Dominik Kahr, Florian Haage, Marietta Zille

**Affiliations:** Department of Pharmaceutical Sciences, Division of Pharmacognosy, University of Vienna, Vienna, Austria; Section of Biomedical Sciences, Department of Health Sciences, Hochschule Campus Wien (HCW) University of Applied Sciences Campus Wien, Vienna, Austria; Department of Pharmaceutical Sciences, Division of Pharmacology and Toxicology, University of Vienna, Vienna, Austria; Vienna Doctoral School of Pharmaceutical, Nutritional and Sport Sciences, University of Vienna, Vienna, Austria

**Author notes:** Corresponding authors: Contact: Ammar Tahir Marietta Zille. Authors contributed equally (Ammar Tahir, Hari Baskar Balasubramanian).

**Keywords:** Doxorubicin, lipidomics, metabolomics, senescence-associated secretory phenotype, tight junctions, untargeted mass spectrometry

## Abstract

The blood-brain barrier (BBB) declines with age, with endothelial senescence implicated as a contributor, yet its metabolic features remain incompletely defined. We induced senescence in mouse brain endothelial cells (BECs) using the chemotherapeutic doxorubicin and the oxidative stress mediator hydrogen peroxide and performed untargeted mass spectrometry of cell lysates and supernatants. Senescence was confirmed by increased senescence-associated β-galactosidase activity and elevated cyclin-dependent kinase inhibitors p16 and p21, along with increased *Il-6* mRNA as senescence-associated secretory phenotype marker. At the BBB structural level, occludin was consistently reduced, ZO-1 decreased with H_2_O_2_ (with a similar trend under doxorubicin), and claudin-5 showed a variable, non-significant reduction, indicating target- and inducer-specific junctional remodeling. Pathway mapping highlighted prominent remodeling of lipid metabolism (including glycerophospholipids, sphingolipids, acylcarnitines) and amino-acid pathways, with both shared and inducer-specific shifts. These discovery-stage *in vitro* results recapitulate several metabolites previously associated with senescence across tissues and nominate additional candidates for hypothesis testing. To increase mechanistic and translational relevance, we outline a staged roadmap: validation in advanced multicellular and flow-based BBB models followed by *in vivo* studies, with attention to route-specific BBB readouts. Overall, this work provides a curated snapshot of metabolic remodeling in brain endothelial senescence that co-occurs with selective tight-junction alterations relevant to BBB aging.

**Graphical abstract:** 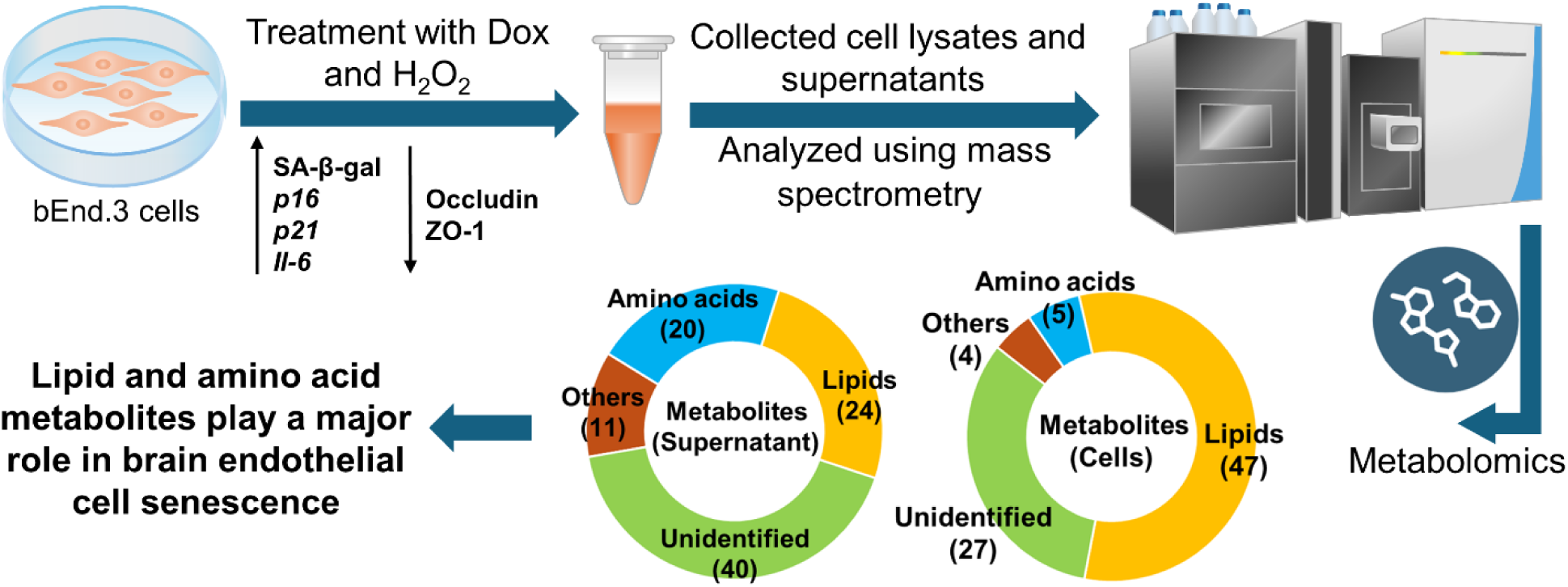

Mouse brain endothelial cells exposed to doxorubicin (Dox) or hydrogen peroxide (H₂O₂) underwent senescence (↑SA-β-gal activity, p16/p21 expression), accompanied by selective reductions in tight-junction proteins. Untargeted mass spectrometry profiled metabolites in cell lysates and supernatants. Lipids, followed by amino acid metabolites, comprised the largest share of alterations, highlighting these pathways as major components of metabolic reprogramming associated in endothelial senescence.

## Introduction

The blood-brain barrier (BBB) separates the peripheral circulation from the brain and restricts the entry of molecules from the bloodstream into the brain (Ballabh et al. 2004). Brain endothelial cells (BECs) line the inner lumen of blood vessels in the BBB and express tight junction proteins, which prevent the leakage of molecules between the cells into the brain, as well as efflux transporters that actively pump molecules out of the brain (Banks et al. 2021; Knox et al. 2022). BECs and the BBB are therefore crucial for maintaining a proper brain homeostasis and their compromise directly affects the onset and progression of cerebrovascular and neurodegenerative diseases, such as stroke and dementia (Archie et al. 2021).

Aging is a phenomenon that naturally occurs in all cell types in our body. Cellular senescence is one of the hallmarks of aging. It is characterized by a state of permanent growth arrest in which cells evade cell death, but remain metabolically active. Cellular senescence is a protective mechanism that halts the proliferation of cells with damaged DNA, thereby preventing tumor formation (Roninson 2003). Beyond its role in cancer suppression, senescence contributes to wound healing and tissue remodeling by secreting signaling molecules that recruit immune cells to support tissue repair and regeneration (Campisi 2005; Muñoz-Espín and Serrano 2014). In mice, senescence has been shown to play a vital role in embryonic organ development by eliminating excess cells and shaping tissue structure (Yao et al. 2023). Additionally, senescent pancreatic beta cells can enhance insulin secretion, highlighting the diverse physiological roles of senescence (Helman et al. 2016). However, the accumulation of senescent cells over time is associated with tissue dysfunction, inflammation, and the progression of age-related pathologies (López-Otín et al. 2023).

During aging, several changes to the morphology and integrity of the BBB occur, such as increased non-specific protein leakage and decreased efflux transporter systems expressed in BECs, pericyte loss, and increased reactivity of astrocytes (Banks et al. 2021; Real et al. 2024). The number of BECs declines with age in humans and the mitochondrial content in BECs decreases in aged monkeys and rats (Erdő et al. 2017). Furthermore, an increase in the number of BECs with high levels of senescence-related gene expression has been observed in a transcriptomics study in aged mice (Kiss et al. 2020). However, a detailed characterization of the senescent phenotype of BECs is currently lacking.

Metabolomics and lipidomics are very useful tools for comprehensive profiling of cellular metabolism. While metabolomics focuses on profiling small molecules, including sugars, amino acids, and organic acids, lipidomics encompasses the profiling of a diverse array of molecules such as fatty acids, phospholipids, and sterols that serve as essential structural components of cell membranes and signaling molecules (Johnson et al. 2016; Han and Gross 2022). Combining metabolite and lipid profiles of cells provides valuable insights into the underlying biochemical pathways and molecular mechanisms (Wang et al. 2020). In the context of aging and neurovascular senescence, metabolomics and lipidomics are now considered promising tools for unraveling the underlying molecular mechanisms responsible for age-related changes in BECs (Panyard et al. 2022; Fang et al. 2023). Recent metabolomics studies showed changes in metabolic pathways associated with aging, such as: changes in glycolysis, AMP/ATP, tricarboxylic acid cycle intermediates (Panyard et al. 2022; Castro et al. 2022), and amino acid metabolism (Sabbatinelli et al. 2019), suggesting metabolic rewiring as a hallmark of cellular senescence. Similarly, lipidome profiling has identified age-related dysregulation of lipid metabolism in aged endothelial cells, characterized by alterations in lipid classes, fatty acid composition, and lipid signaling molecules (Sharma and Diwan 2023; Giuliani et al. 2023).

In this study, we sought to identify the metabolites associated with senescence in BECs. For this, we established an *in vitro* model of BEC senescence, using the well-known senescence inducer doxorubicin (Dox) (Marques et al. 2020), and the pro-oxidant hydrogen peroxide (H_2_O_2_). Using the two different inducers, we identified common metabolites likely to be related to senescence using untargeted metabolomics and lipidomics (**Fig. S1**).

Here, we present a discovery-stage untargeted metabolomic and lipidomic profiling of brain endothelial cell senescence paired with tight-junction readouts, to prioritize candidate pathways and metabolites for follow-up. We explicitly outline a staged translational roadmap to increase mechanistic resolution and relevance, from monoculture discovery to validation in advanced multicellular and flow-based BBB models (Bhalerao et al. 2020; Chaulagain et al. 2023), followed by *in vivo* confirmation. This approach complements *in vivo* evidence that endothelial senescence contributes to BBB aging and neurovascular decline (Kiss et al. 2020; Knopp et al. 2023; Novo et al. 2024).

## Methods

### Cell culture

The mouse BEC cell line bEnd.3 (ATCC #CRL-2299) up to passage 11 was cultivated in Dulbecco’s modified Eagle’s medium (ThermoFisher #31966047) supplemented with 10 % fetal bovine serum (Sigma #F7524) and 1 % penicillin/streptomycin (Sigma #P0781) at 37 °C and 5 % CO_2_. They were harvested with accutase (Sigma #A6964) and seeded at a density of 50 000 cells/well on coverslips in 24-well plates for SA-β-gal staining, 60 000 cells/well in 12-well plates for gene expression, or 150 000 cells/well in 6-well plates for immunoblotting. The next day, the cells were incubated with 100 nM Dox (Sigma #D1515) or 750 µM H_2_O_2_ (Sigma #14533) for 72 hours.

### Senescence-associated β-galactosidase assay

The assay was performed as previously described (Debacq-Chainiaux et al. 2009) with modifications. The cells were fixed in 2 % formaldehyde (Sigma #252549) and 0.2 % glutaraldehyde (Sigma #3802) in phosphate-buffered saline (PBS, Gibco #18912-014) for 3 minutes. After washing twice with PBS, the cells were stained with freshly prepared staining solution containing 150 mM sodium chloride (Sigma #S9888), 40 mM citric acid (Sigma #C0759)/sodium phosphate buffer (Sigma #S5136), 5 mM potassium hexacyanoferrate (III) (Sigma #244023), 5 mM potassium hexacyanoferrate (II) trihydrate (Sigma #P9387), 2 mM magnesium chloride (Sigma #M0250), and 1 mg/ml 5-bromo-4-chloro-3-indolyl β D-galactopyranoside (X-gal, Sigma #B4252) dissolved in N,N-dimethylformamide (Sigma #270547) for 24 hours at 72 °C under CO_2_ exclusion. Thereafter, the cells were washed twice with PBS and nuclear staining was performed using Hoechst 33342 (Sigma #14533, 6 μg/ml in PBS) for 10 minutes. After washing, the coverslips were mounted onto glass slides using fluoromount (ThermoFisher #00-4958-02).

### Microscopy and image analysis

We performed SA-β-gal imaging at an Olympus BX51 microscope with a UC90 color camera. Three random positions on each coverslip were selected based on the Hoechst staining and bright-field images were acquired from the same positions. Image analysis was performed using ImageJ (v.1.53q, RRID:SCR_003070) semi-automatically to minimize the bias of manual counting. We used a color threshold (Hue 91-170, Saturation 30-255, Brightness 77-149) to detect the SA-β-gal-positive cells and counted the number of particles using the “Analyze particles” function (size threshold 60-infinity, as determined in comparison to manual counting). The total number of cells per Hoechst image was determined using the “Threshold” function, removing background noise with “despeckle”, applying “watershed” to separate the nuclei, and finally using “Analyze particles” with no size threshold. The sum percentage was determined by adding all SA-β-gal-positive cells divided by the sum of all Hoechst-positive cells per condition to minimize bias of individual images.

### RNA extraction and real-time PCR

We prepared total RNA using E.Z.N.A. HP Total RNA Kit (VWR #R6812) according to the manufacturer’s protocol. RNA concentration and purity were determined spectrophotometrically (NanoDrop 2000, ThermoScientific) by assessing A260/A280 and A260/A230 ratios. For each sample, 10-15 ng of total RNA was used per reaction. Real-time PCR was performed using VWR® Probe One-Step RT-qPCR Master Mix with UDG (VWR #73224). TaqMan assays (ThermoFisher) were used for mouse *P16/Cdkn2a* (#Mm00494449_m1), *P21/Cdkn1a* (#Mm00432448_m1), *Il-6* (#Mm00446190_m1), and the endogenous control *b-actin* (#Mm02619580_g1). Amplification was performed on a Thermo ABI QuantStudio 5 real-time PCR system with the following cycling conditions: 25 °C for 30 s, 55 °C for 10 min, 95 °C for 1 min, followed by 45 cycles of 95 °C for 10 s and 60 °C for 1 min. All reactions were performed in technical triplicates, with no-template and no-reverse transcription controls included to rule out contamination or genomic DNA amplification. Primer/probe sets were pre-validated by the manufacturer. Assay efficiencies were determined from standard curves and fell within 90-110%. Relative expression levels were calculated using the ΔΔCt method with β-actin as reference. Six independent biological replicates were analyzed.

### Immunoblot analysis (Western Blotting)

We prepared protein extracts using 1x RIPA buffer (ThermoFisher #89900) supplemented with protease inhibitor cocktail (cOmplete™, Mini, EDTA-free Protease-Inhibitor-Cocktail, Sigma #11836170001). Lysates were incubated on ice and clarified by centrifugation at 21000xg for 10 min, and supernatants were collected. Protein concentrations were determined using the Bradford assay (Quick Start™ Bradford 1x Dye Reagent, Bio-rad #5000205) and equal amounts of protein (15 µg per lane) were mixed with Laemmli sample buffer (ROTI®Load 2 (Gel loading Laemmli buffer, 4X, non-reducing, Roth #H318) containing β-mercaptoethanol (Sigma #63689), denatured at 95 °C for 5 min, and subjected to SDS-PAGE under reducing conditions on 6 % (ZO-1 and vinculin), 12 % (P21 and occludin), and 15 % (claudin) polyacrylamide gels.

Electrophoresis was performed at 100 V for 10 min (stacking gel), followed by 150-200 V for 50-60 min (resolving gel). Proteins were transferred onto nitrocellulose membranes (0.2 µm pore size, Amersham™ Protran®, #GE10600001) semi-dry using the Trans-Blot Turbo Transfer System (Bio-rad #704150) at 25 V for 30 min. Membranes were blocked in 5 % bovine serum albumin fraction V (Roth #80764) in Tris-buffered saline with 0.1 % Tween-20 (TBST, 20 mM Tris base, Sigma #T6066; 150 mM NaCl, Sigma #3014; 0.1 % Tween-20, Sigma #P1379) for 1 h at room temperature, followed by overnight incubation at 4 °C with primary antibodies against claudin-5, occludin, ZO-1, p21, and vinculin (see **Supplementary Table 1** for details). After washing in TBST (3× 5 min), membranes were incubated with appropriate HRP-conjugated secondary antibodies (see **Supplementary Table 2** for details) for 1 h at room temperature.

Protein bands were visualized using enhanced chemiluminescence (SuperSignal^TM^ West Femto, ThermoFisher #34094) and imaged with the ChemiDoc MP Imaging System (Bio-rad). Densitometric analysis was performed using ImageJ 1.54g, and protein expression levels were normalized to vinculin as loading control. To note, we did not use GAPDH, although proteins on 12 and 15 % gels were probed for it, because GAPDH increased under H_2_O_2_ conditions. Therefore, vinculin had to be used for all target proteins. We ensured maximal comparability by preparing and loading of each sample per biological replicate in parallel on the different gels.

### Sample preparation for metabolomics analysis

Cells were counted manually using the plugin “Cell Counter” in ImageJ to ensure consistency across samples and to, later on, normalize the metabolite intensities to the cell count. The supernatant media containing metabolites were collected, three volumes of 100 % pre-cooled methanol (VWR #83638) were added, and the lysates were incubated at −80 °C for a minimum of 2 hours. The cells were washed with ice-cold PBS and incubated in a pre-cooled 80% methanol solution for 5 minutes. The cells were scraped, and the samples were then incubated at −80°C for a minimum of 2 hours, to facilitate metabolite extraction. After incubation, both media and cell samples were centrifuged at maximum speed for 15 minutes at 4°C to separate cellular debris, and the supernatants were carefully removed and transferred to new tubes. The supernatants were then evaporated in a SpeedVac apparatus at 45°C for approximately 3-4 hours to concentrate the metabolite extract. The dried samples were reconstituted in 80 % methanol solution at room temperature, followed by vortexing for 10 seconds and a 5-minute ultrasonication bath. The samples underwent a 10-minute centrifugation step at maximum speed. Finally, clarified samples were transferred to LC-MS vials for subsequent metabolomics analysis. Quality control (QC) samples were prepared by collecting from each sample.

### Assay statistical analysis

For SA-β-gal analyses, gene expression and immunoblot analysis, Kruskal Wallis test followed by targeted Mann-Whitney U posthoc tests were performed. Bonferroni-Holm correction was used to adjust for the inflation of Type I error because of multiple testing. Data are presented as medians. A value of *P* < 0.05 was considered statistically significant.

### Metabolomics: Separation and detection

Samples were separated using an Exion UHPLC system (Sciex, Darmstadt Germany) using a RP-C18 Kinetex 2.1×100 mm, 1.7 µm, 100 Å column (Phenomenex, Germany) using a binary 5-minute gradient: Solvent A (5 % acetonitrile) and solvent B (95 % acetonitrile), both buffered with 5 mM ammonium formate. The gradient details were: 0-0.5 min = 100 % A, 3.5 min = 100 % B, 3.6-5 min =100 % A. The injection loop, needle, and seat were washed after each injection with a rinse solution of methanol, 2-propanol and water (40:40:10, vol%/vol%) to ensure no carry over or contamination happened. HRMS detection was performed on a X-500 QTOF system (Sciex, Darmstadt, Germany). The detection setup comprised of an independent data acquisition with top 10 most abundant ions experiment with the following parameters: Range = 50-1500 Da, gas1 = 30 psi, gas2 = 35 psi, curtain gas = 45 psi, spray voltages = 4.5 KV/+5.0 KV for negative and positive ionization, collision activated dissociation fragmentation = Dynamic 35 V (−/+ 15 V) collision energy, accumulation time = 0.1 ms and decluttering potential = −80 V.

Lipids and metabolites identification and statistics was performed according to our previously established pipeline (Tahir et al. 2024). Briefly, raw data were processed using MS-DIAL (v4.9.221218, Windows x64) (Tsugawa et al. 2020), applying the following key settings: minimum peak height of 1000, mass slice width 0.05 Da, MS2Dec sigma window 0.5, MS/MS abundance cutoff 10, and mass tolerance for identification and alignment set at 0.005 Da. Retention time alignment used a tolerance of 0.05 min.

### Metabolomics: Validation and normalization

For the analytical validation of our workflow, we first assessed the level of analytical variation across the dataset. This initial evaluation was performed prior to applying any QC-based LOESS normalization or batch correction, in order to examine the raw data quality and consistency. Specifically, we inspected the clustering of QC samples and assessed signal drift and intensity variation across runs. This step allowed us to verify that the observed variation was primarily technical rather than biological, and to justify the need for subsequent normalization. Only after confirming this, we applied the appropriate correction strategies to minimize batch effects and ensure comparability across all sample groups. The dataset was normalized and batch-corrected using LOESS (Lai et al. 2018) and internal standard-based workflows. High-quality features were selected based on 2-way ANOVA p-values and RSD filtering.

### Metabolomics: Identification

Lipid identification utilized LipidBlast (Kind et al. 2013) in both ion modes, with all relevant adducts and modifiers selected. Metabolites were identified using the ESI(+)-MS/MS database (Kind et al. 2013), and further matched against HMDB, METLIN Gen2 and other databases when necessary. Final curation involved manual inspection of MS2 spectra (dot product score >0.75), duplicate removal, and adduct validation. When MS2-based identification was not feasible using the integrated spectral libraries, we performed HRMS putative peak annotation using external databases including HMDB (Wishart et al. 2007), METLIN Gen2 (acquired on 20.01.2023), and the Metabolomics Workbench (https://www.metabolomicsworkbench.org/), applying a mass accuracy threshold of ±5 ppm. The **Supplemental data file** includes the list of identified metabolites. Metabolites marked with an asterisk (*) indicate putative annotations based on spectral matching without confirmation by reference standards. All other metabolites were identified with higher confidence by matching both MS¹ and MS² spectra against curated libraries.

In line with our commitment to transparency and data sharing, we have deposited all relevant data in the MassIVE repository (as detailed in the Data Availability section). The repository contains the original Sciex raw files, the full MS-DIAL project, and the uncurated metabolite identification tables exported from MS-DIAL, including normalized values. The repository can be accessed via the following link: https://doi.org/doi:10.25345/C5QJ78B1G

### Metabolomics: Statistics and Interpretation

Furthermore, the dataset of identified features was further analyzed using MetaboAnalyst (Xia and Wishart 2010a; Xia and Wishart 2010b; Pang et al. 2021). Initial significance testing was performed through volcano plot analysis to highlight differentially abundant features across conditions. To assess the overall structure and quality of the dataset, PLSDA was applied to the three experimental groups (Veh, H_2_O_2_, and DOX), allowing us to evaluate the distinct clustering and separation between cohorts. In addition, we used hierarchical clustering and heatmap visualization to examine global patterns in feature abundance and group-specific expression profiles. Finally, metabolites showing significant regulation were submitted to the Reactome Pathway Browser (Milacic et al. 2024) (https://reactome.org/PathwayBrowser/) and PathWhiz (https://www.smpdb.ca/pathwhiz/) for functional and qualitative pathway mapping.

## Results

### Senescence validation and tight-junction changes in BECs

First, we established an *in vitro* model of BEC senescence. The induction of cellular senescence has been most commonly and reliably achieved using chemical agents such as chemotherapeutic drugs and oxidizing compounds. The chemotherapeutic drug Dox disrupts DNA by intercalating into its structure, while oxidizing agents, such as H_2_O_2_, induce oxidative stress and thereby cause DNA damage. Both agents activate the DNA damage response pathway (Kudlova et al. 2022). However, data on BEC senescence induction are limited.

Therefore, we treated bEnd.3 cells with Dox for 72 hours at different concentrations (5-500 nM) to determine the optimal concentration for senescence induction (**Fig. S2**). The number of senescence-associated β-galactosidase (SA-β-gal)-positive cells increased from 25 nM and was significantly elevated at 100 nM, whereas the number of Hoechst-positive cells, as a measure of the total cell count, decreased. Starting from 250 nM, we observed cell death in some cultures, which was even more prominent at 500 nM. Therefore, we decided to continue experiments with 100 nM Dox (**Fig. 1a-b**).

**Figure 1:**
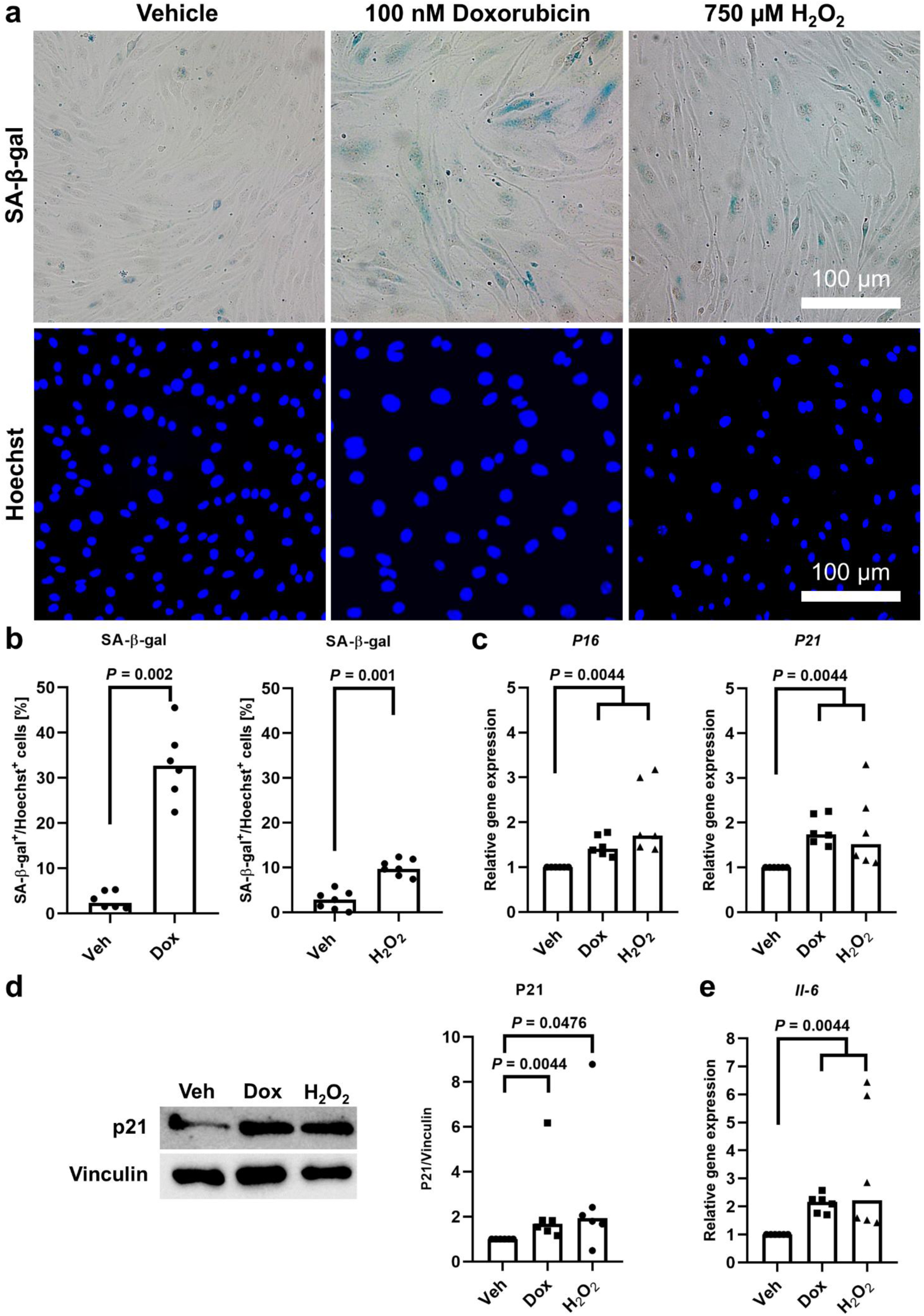
Senescence of BECs induced by doxorubicin and hydrogen peroxide. Senescence was induced in BECs by incubation with 100 nM Dox or 750 µM H_2_O_2_ for 72 hours. **a-b)** The percentage of SA-β-gal-positive cells significantly increased after Dox (N = 6/group) and H_2_O_2_ (N = 7/group) treatment. Mann-Whitney U test. **c)** *P16* and *P21* mRNA expression relative to β-actin increased in Dox- and H_2_O_2_-treated cells (N = 6/group). Kruskal Wallis test: *χ^2^*(2, N = 18) = 12.78, *P* < 0.0001, *η²* = 0.752, followed by targeted Mann-Whitney U tests with Bonferroni-Holm correction. **d)** P21 protein levels increased in Dox- and H_2_O_2_-treated cells (N = 6/group). Protein levels were normalized to vinculin loading control that does not change in response to Dox or H_2_O_2_. Kruskal Wallis test: *χ^2^*(2, N = 18) = 8.20, *P* = 0.0107, *η²* = 0.482, followed by targeted Mann-Whitney U tests with Bonferroni-Holm correction. **e)** SASP marker *Il-6* mRNA expression increased in Dox- and H_2_O_2_-treated cells (N = 6/group). Kruskal Wallis test: *χ^2^*(2, N = 18) = 11.79, *P* = 0.0005, *η²* = 0.694, followed by targeted Mann-Whitney U tests with Bonferroni-Holm correction.

To generate a model of BEC senescence more relevant to cerebrovascular and neurodegenerative diseases, we next examined H_2_O_2_ (25-1000 µM for 72 hours) as a potential inducer (**Fig. S3**). The concentration-response demonstrated an increase in the number of SA-β-gal-positive cells and a decrease in the number of Hoechst-positive cells starting at 250 µM, with a significant difference compared to vehicle (Veh) treatment at 750 µM and noticeable cell death at 1000 µM. Therefore, we decided to continue experiments with 750 µM H_2_O_2_ (**Fig. 1a-b**).

Next, we validated the senescence phenotype of both Dox- and H_2_O_2_-treated BECs using the cell cycle regulatory proteins p16^INK4a^ (p16) and p21^WAF1^ (p21). Both treatments increased *p16* and *p21* gene expression relative to Veh (**Fig. 1c**) as well as p21 protein expression (**Fig. 1d**). In addition, both showed increased gene expression of the senescence-associated secretory phenotype (SASP) marker *Il-6* (**Fig. 1e**).

To assess barrier-related structural changes, we measured the abundance of claudin-5, occludin, and ZO-1 by immunoblot. Both Dox and H_2_O_2_ consistently reduced occludin protein levels relative to vehicle (**Fig. 2**). ZO-1 was significantly decreased with H_2_O_2_ and showed a similar but non-significant decrease with Dox. Claudin-5 exhibited higher variability and did not reach statistical significance. These data indicate inducer- and target-specific junctional remodeling, with occludin most robustly affected under our conditions.

**Figure 2:**
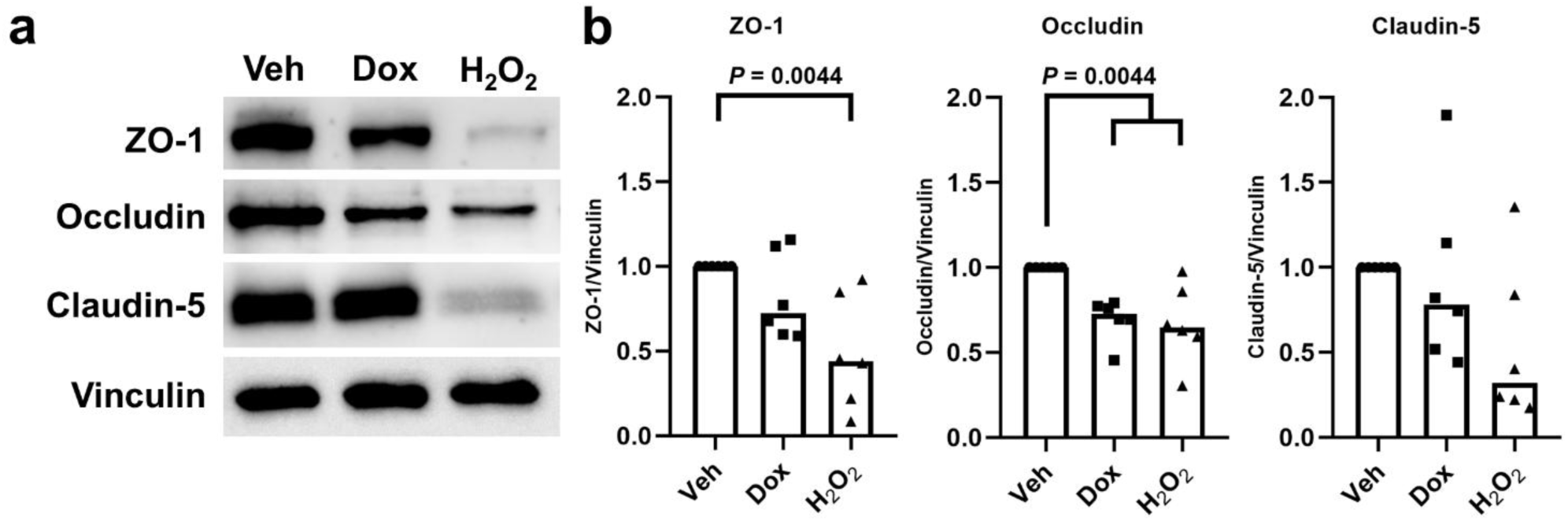
Tight junction proteins are decreased in senescent BECs. Senescence was induced in BECs by incubation with 100 nM Dox or 750 µM H_2_O_2_ for 72 hours. **a)** Representative immunoblots of the levels of the tight junction proteins ZO-1, occludin, claudin-5 and vinculin loading control that does not change in response to Dox or H_2_O_2_. **b)** Tight junction levels relative to vinculin decreased after H_2_O_2_ treatment for ZO-1 and occludin and after Dox treatment only for occludin, while claudin-5 showed higher variability (N = 6/group). Kruskal Wallis test followed by targeted Mann-Whitney U tests with Bonferroni-Holm correction: ZO-1 *χ^2^*(2, N = 18) = 8.35, *P* = 0.0086, *η²* = 0.491; Occludin *χ^2^*(2, N = 18) = 11.90, *P* = 0.0004, *η²* = 0.700; Claudin-5 *χ^2^*(2, N = 18) = 4.70, *P* = 0.0919, *η²* = 0.276. Uncropped blots with molecular weight markers are provided in **Fig. S4**.

### Discovery-stage metabolomic and lipidomic remodeling in senescent BECs

After successfully inducing senescence in our cell culture model, we sought to investigate the molecular profile of both treated cells (Dox and H_2_O_2_) compared to the Veh-treated control. To do this, we extracted both the cells and the supernatants, which were then subjected to untargeted lipidomics and metabolomics. Partial Least Squares Discriminant Analysis (PLSDA) analyses illustrated the distribution of three groups, Dox-treated, H_2_O_2_-treated, and Veh samples (**Fig. 3a and 4a**). Although there was a distinct separation of all groups, the Dox and H_2_O_2_ clusters were closer together compared to the Veh cluster, suggesting commonalities between their metabolic profiles, which may be those metabolites related to senescence.

**Figure 3:**
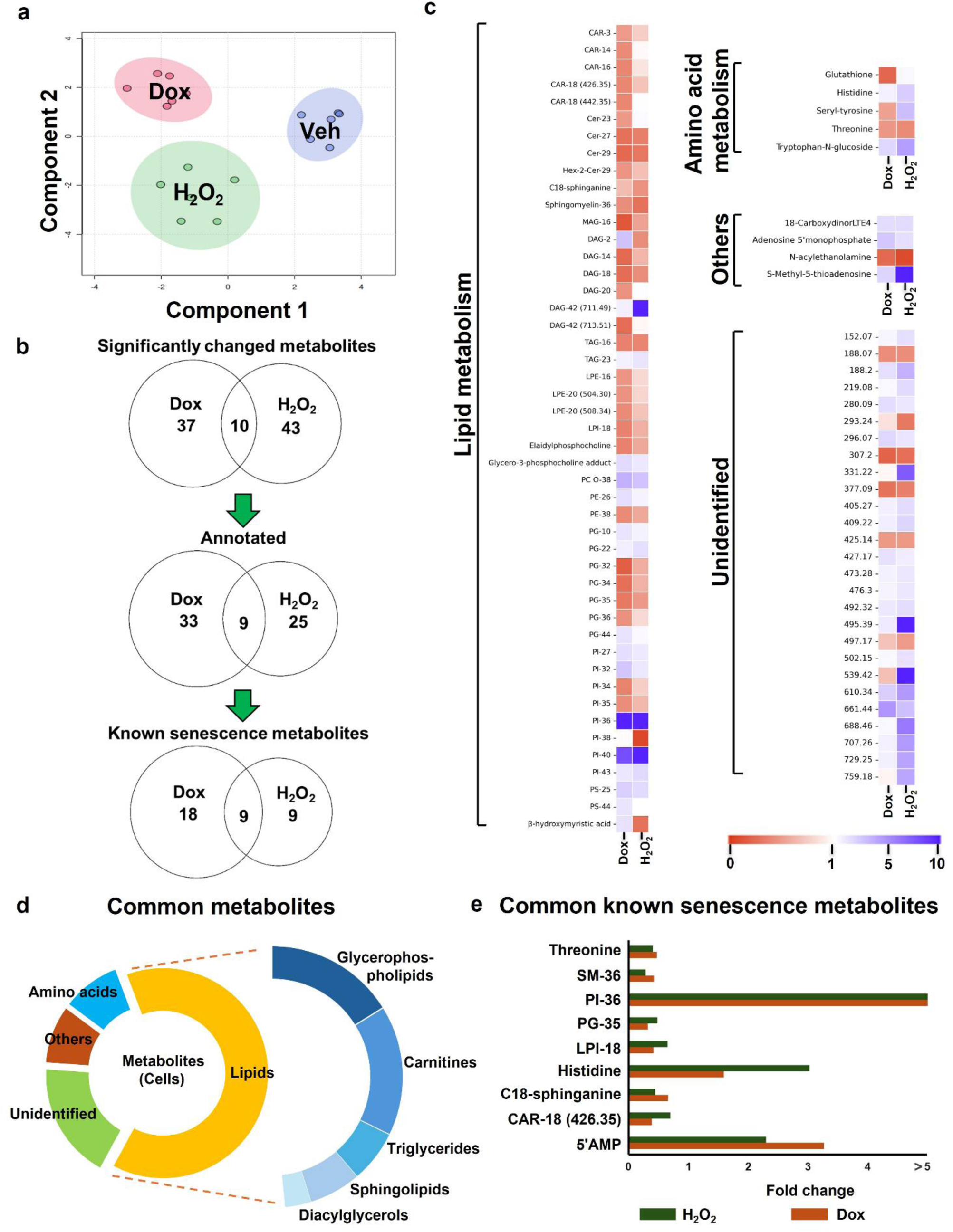
Metabolomic pattern associated with BEC senescence in cells. **a)** PLSDA score plot of both inducers of senescence (Dox and H_2_O_2_) versus vehicle. **b)** Venn diagrams represent the significantly changed metabolites (threshold fold-change ≥ 2 for increased or ≤ 0.5 for decreased metabolites). **c)** Heatmap of metabolites fold changes between Dox/H_2_O_2_ and vehicle. Heatmaps were generated in Jupyter notebook using Python(V3). Data were preprocessed and formatted using Pandas function. Heatmaps were generated with Seaborn, and figure customization and rendering were performed using Matplotlib function. The number between the brackets indicates the m/z in Da of the metabolites. **d)** Stacked donut chart of metabolite groups and subclasses. Metabolites classified as others neither belong to lipids nor amino acids. **e)** Fold-change of known senescence metabolites common between Dox and H_2_O_2_. 5’AMP = adenosine 5’monophosphate, CAR = carnitine, Cer = ceramide, DAG = diacylglycerides, Hex = hexanoyl, LPE = lysophosphatidylethanolamine, LPI = lysophosphatidylinositol, MAG = monoacylglycerides, PE = phosphatidylethanolamine, PG = phosphaditylglycerol, PI = phosphatidylinositol, PS = phosphaditylserine, PC O = ether-linked phosphatidylcholine, TAG = triacylglycerides.

**Figure 4:**
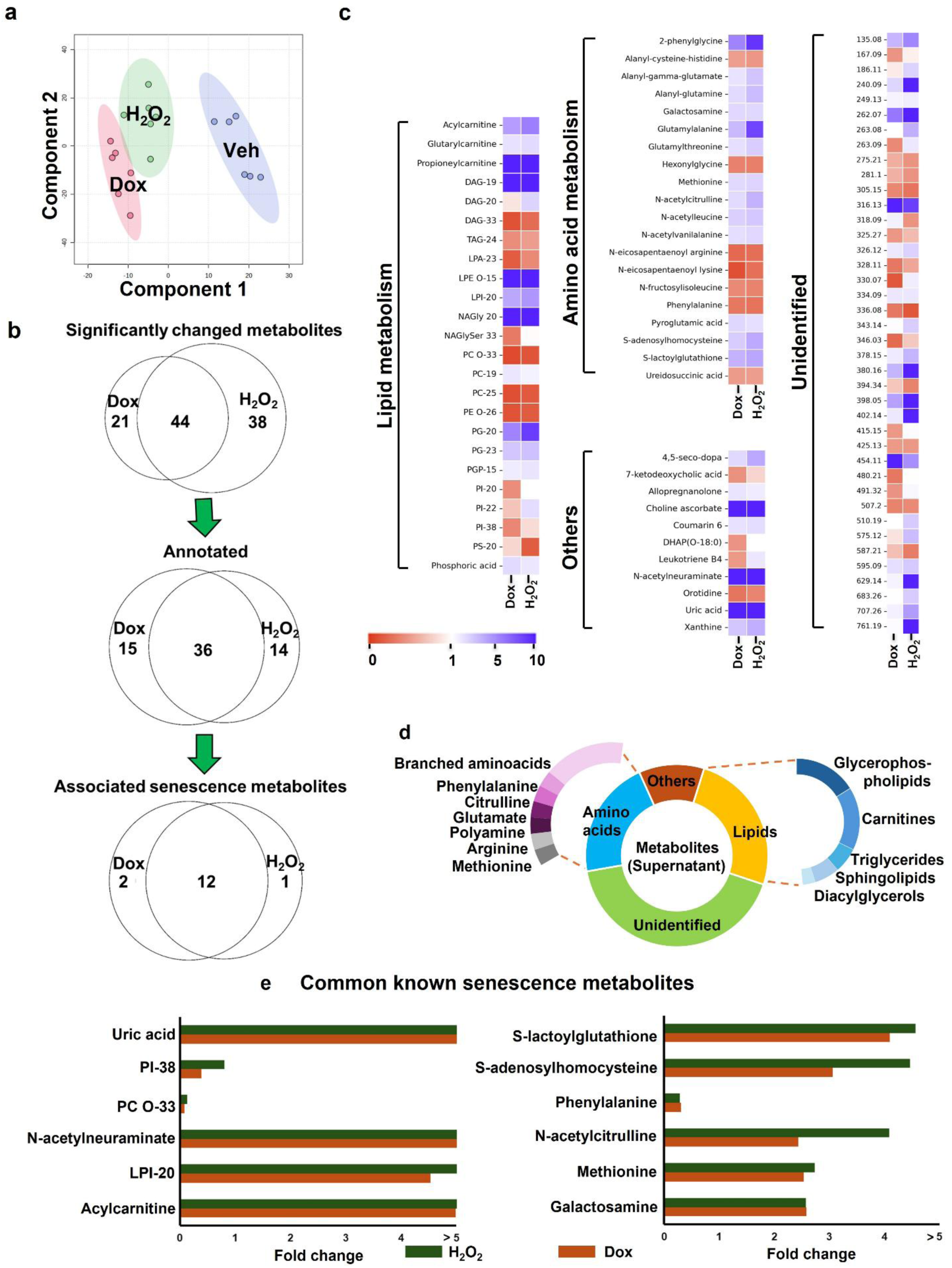
Metabolomic pattern associated with BEC senescence in supernatant samples. **a)** PLSDA score plot of both inducers of senescence (Dox and H_2_O_2_) versus vehicle. **b)** Venn diagrams represent the significantly changed metabolites (threshold fold-change ≥ 2 for increased or ≤ 0.5 for decreased metabolites). **c)** Heatmap of metabolite fold changes between Dox/H_2_O_2_ and vehicle. Heatmaps were generated in Jupyter notebook using Python(V3). Data were preprocessed and formatted using Pandas function. Heatmaps were generated with Seaborn, and figure customization and rendering were performed using Matplotlib function. The number between the brackets indicates the m/z in Da of the metabolites. **d)** Stacked donut chart representation of metabolite groups and subclasses. Metabolites classified as others neither belong to lipid nor amino acids. **e)** Fold-change of known senescence metabolites common between Dox and H_2_O_2_. CAR = carnitine, DAG = diacylglycerides, DHAP = dihydroxyacetone phosphate, LPA = lysophosphatidic acid, LPE = lysophosphatidylethanolamine, LPI = lysophosphatidylinositol, NAGly =N-acyl glycine, NAGlyser = N-acyl-glycyl serine, PE = phosphatidylethanolamine, PE O = ether-linked PE, PC = phosphatidylcholine, PC O = ether-linked PC, PG = phosphaditylglycerol, PGP = phosphatidylglycerophosphate, PI = phosphatidylinositol, PS = phosphaditylserine, TAG = triacylglycerides.

We identified 47 metabolites with significant differential changes (*P* < 0.05) within cells and 65 in the supernatant of Dox-induced senescent cells (**Fig. 3b and 4b**, **Supplemental data file**). In comparison, 53 metabolites showed differential changes in cells and 82 in the supernatant of H_2_O_2_-induced senescent cells. Among these, 10 metabolites were commonly altered within the cells, and 44 were common in the supernatant across both treatment conditions.

In the cell lysates, we were able to identify 42 metabolites in the Dox-treated group and 34 in the H_2_O_2_ group, of which 9 metabolites overlapped (**Fig. 3b**, **Supplemental data file**). The majority of the identified metabolites were associated with fatty acid metabolism (47), and only a few amino acid (5) and other metabolites (4) were changed (**Fig. 3c-d, Supplemental data file**). Lipid metabolites such as ether-linked-phosphatidylcholine (PC O)-38, phosphatidylinositol (PI)-36 and −40, and phosphatidylserine (PS)-25 were increased (fold-change > 2) in both H_2_O_2_ and Dox treatments compared to vehicle, whereas ceramide-27 and −29, diacylglyceride (DAG)-18, phosphatidylglycerol (PG)-35, sphingomyelin-36, and triglyceride (TAG)-16 were decreased (fold-change < 0.5). In addition, some metabolites were changed only in the Dox group (e.g., carnitines) or only in the H_2_O_2_ group (e.g., PI-38). Among the amino acid metabolites, tryptophan-N-glucoside was increased in both treatments compared to vehicle, whereas threonine was decreased. Other metabolites increased in both treatments were 18-carboxydinorLTE4, 5-methyl-5-thioadenosine, and adenosine 5’-monophosphate, whereas N-acylethanolamine was decreased.

In the supernatant, we identified 51 metabolites in the Dox group and 50 in the H_2_O_2_ group, of which 36 metabolites overlapped (**Fig. 4b**, **Supplemental data file**). Compared to the cell lysates, the number of lipid metabolites (24) was similar to that of amino acid metabolites (20), followed by other metabolites (11) (**Fig. 4c-d, Supplemental data file**). Among lipid metabolites, CAR (acylcarnitine, glutarylcarnitine, propenoylcarnitine), DAG-19, phosphoric acid, ether-linked-phosphaditylethanolamine (LPE O-15), lysophosphatidylinositol (LPI)-20, N-acyl glycine (NAGly)-20, and phosphaditylglycerol (PG)-20 and −23 were increased in H_2_O_2_ and Dox treatments compared to vehicle, whereas DAG-33, lysophosphatidic acid (LPA)-23, phosphatidylcholine (PC)-25, ether-linked-phosphatidylethanol (PE O)-26, and ether-linked-phosphatidylcholine (PC O)-33 were decreased. In addition, some metabolites were changed only in the Dox group (e.g., PI-38 and N-acyl glycyl serine (NAGlySer)-33) or only in the H_2_O_2_ group (e.g., PS-20). Among amino acid metabolites, 2-phenylglycine, alanyl-gamma-glutamate, alanyl-glutamine, galactosamine, glutamylalanine, glutamylthreonine, methionine, N-acetylcitrulline, N-acetyl leucine, N-acetylvanilalanine, pyroglutamic acid, S-adenosylhomocysteine, and S-lactoylglutathione were increased in both treatments, whereas hexonylglycine, N-eicosapentaenoyl arginine, N-eicosapentaenoyl lysine, N-fructosylisoleucine, and phenylalanine were decreased.

### Metabolic signatures of senescence in BECs

To evaluate the relevance of our findings, we compared metabolites consistently altered by both senescence inducers (Dox and H_2_O_2_) with those reported in well-characterized senescence models and human aging cohorts (**Table 1**).

**Table 1.** Comparison of identified metabolites with senescence/aging metabolites from the literature. Directionality refers to changes in senescent versus control conditions. Differences between studies may reflect variation in cell type, senescence trigger, or *in vivo* versus *in vitro* context. Cmah = cytidine monophosphate-N-acetylneuraminic acid hydroxylase, Dox = doxorubicin, LPI = lysophosphatidylinositol, PI = phosphatidylinositol, PG = phosphatidylglycerol, PC O = (ether-linked-phosphatidylcholine), SAMP8 = senescence-accelerated mouse-prone 8.

| Metabolite | System (literature) | Senescence model (literature) | Refer<br>ence | Direction (literature) | Direction (detected in this study in) |
| --- | --- | --- | --- | --- | --- |
| Lipid metabolism |  |  |  |  |  |
| C2-carnitine (Acylcarnitine) | Human serum | Aging cohort (SASP-high individuals) | (Ham sanat han et al. 2024) | ↓ | ↓ (supernatant) |
| Carnitine-18 |  |  |  | ↓ | ↓ (cells) |
| Sphingomyelin-36 |  |  |  | ↓ | ↓ (cells) |
| Sphinganine | Human oral fibroblast cell line (NHOF-1) | Replicative senescence | (Jame s et al. 2016) | ↑ | ↓ (cells) |
| LPI-18 | Human MCF-7 breast cancer cells | Oncogene-induced senescence (NeuT) | (Cade nas et al. 2012) | ↑ | ↓ (cells) |
| PG-35 |  |  |  | ↑ | ↓ (cells) |
| PI-36 |  |  |  | ↑ | ↑ (cells) |
| LPI-20 |  |  |  | ↑ | ↑ (supernatant) |
| PC O-33 |  |  |  | ↑ | ↓ (supernatant) |
| Amino acid metabolism |  |  |  |  |  |
| Histidine | Human serum | Aging cohort (SASP-high individuals) | (Ham sanat han et al. 2024) | ↓ (associated with higher CRP / aging) | ↑ (cells) |
|  | Mouse muscle | SAMP8 (accelerated aging) | (Hosh ino et al. 2022) | ↓ |  |
| Threonine | Human serum | Aging cohort (SASP-high individuals) | (Ham sanat han et al.<br>2024) | ↓ (associated with higher CRP / aging) | ↓ (cells) |
|  | Mouse muscle | SAMP8 (accelerated aging) | (Hosh<br>ino et<br>al.<br>2022) | ↓ |  |
| Methionine | Mouse muscle | SAMP8 (accelerated aging) | (Hosh<br>ino et<br>al.<br>2022) | ↑ | ↑ (supernatant) |
| Phenylalanine |  |  |  | ↓ | ↓ (supernatant) |
| N-acetyl citrulline | Human serum | Aging cohort (SASP-high<br>individuals) | (Ham<br>sanat<br>han et<br>al.<br>2024) | ↑ | ↑ (supernatant) |
| S-adenosyl-homocysteine | Rat vascular smooth muscle<br>cells | Metabolite-induced<br>senescence | (Mi et<br>al.<br>2022) | ↑ | ↑ (supernatant) |
|  | Mouse muscle | SAMP8 (accelerated aging) | (Hosh<br>ino et<br>al.<br>2022) | ↑ |  |
| S-lactoylglutathione | Human oral fibroblast cell<br>line (NHOF-1) | Replicative senescence | (Jame<br>s et<br>al.<br>2016) | ↑ | ↑ (supernatant) |
| Galactosamine | Mouse hepatocytes | Chemical (Dox and cisplatin)<br>/ oxidative stress (H<sub>2</sub>O<sub>2</sub>)<br>induced senescence | (Kum<br>ar et<br>al.<br>2024) | ↑ | ↑ (supernatant) |
| **Other metabolic pathways** |  |  |  |  |  |
| Adenosine 5'monophosphate | Mouse muscle | SAMP8 (accelerated aging) | (Hosh<br>ino et al.<br>2022) | ↑ | ↑ (cells) |
| N-acetylneuraminate | Mouse adipose tissue | Cmah knockout (senescence modulation model) | (Take da et al.<br>2023) | inferred ↑<br>(senescence-associated pathway) | ↑ (supernatant) |
| Uric acid | Mouse lens epithelial cells | Hyperuricemia-induced senescence | (Lin et al.<br>2024) | ↑ | ↑ (supernatant) |

Lipid metabolites represented a major overlapping class between our dataset (**Fig. 3e**, **4e**) and the literature, which is consistent with the well-established role of lipid metabolism in regulating cellular senescence (Hamsanathan and Gurkar 2022). For example, we observed decreased levels of long-chain acylcarnitines (e.g., CAR-18 in cell lysates and acylcarnitine in supernatant), in agreement with findings in human serum from individuals with circulating senescence-associated secretome markers, indicative of systemic aging-associated metabolic remodeling (Hamsanathan et al. 2024).

Similarly, sphingolipid alterations were evident across models: sphingomyelin-36 was reduced both in our senescent BECs and in human aging serum (Hamsanathan et al. 2024). In contrast, sphinganine showed model-dependent behavior, being decreased in our cell lysates but previously reported to increase in replicative senescence of human oral fibroblasts (James et al. 2016), highlighting potential cell type- or senescence trigger-specific differences.

Glycerophospholipid remodeling was also observed, with alterations in LPI-18, PG-35, and PI-36 in cell lysates, and LPI-20 and PC O-33 in the supernatant. Comparison with oncogene-induced senescence in MCF-7 human breast cancer cells (NeuT overexpression model) (Cadenas et al. 2012) revealed partial overlap: while PI-36 and LPI-20 showed concordant regulation, LPI-18, PG-35, and PC O-33 displayed opposite directional changes. This suggests that, although membrane lipid remodeling is a conserved feature of senescence, the specific lipid species and their directionality are strongly influenced by cellular context and the mode of senescence induction.

Alterations in amino acid metabolism were also consistently observed across both senescence inducers, with histidine increased and threonine decreased in the cell lysates (**Fig. 3e**), and methionine increased and phenylalanine decreased in the supernatant (**Fig. 4e**). These metabolites have previously been reported in human serum from individuals with circulating senescence-associated secretome markers, reflecting a systemic aging context (Hamsanathan et al. 2024), as well as in skeletal muscle of aged (40- and 55- vs. 12-weeks-old) senescence-accelerated mouse-prone 8 (SAMP8) mice, representing an *in vivo* model of accelerated aging (Hoshino et al. 2022). In these studies, histidine and threonine were reduced in aged or high-inflammatory states, which partially contrasts with our observations in BECs, suggesting cell type-specific regulation. Notably, the increase in methionine and decrease in phenylalanine observed in our dataset is consistent with their elevation and decrease, respectively, in aged SAMP8 mouse muscle [29], indicating conserved alterations in these metabolic pathways across senescence models. Overall, several metabolites showed conserved regulation across models (e.g., acylcarnitines, threonine, methionine), whereas other diverged by cell type and inducer (e.g. sphinganine, histidine), consistent with context-dependent metabolic rewiring in senescence (Fang et al. 2023).

N-acetyl citrulline, which was elevated in our dataset under both treatments, has similarly been reported to increase in the serum of male rapid agers (Hamsanathan et al. 2024), supporting a conserved increase associated with systemic senescence phenotypes. S-adenosyl-homocysteine was also increased following senescence induction in BECs. Notably, this metabolite has been shown to directly induce senescence in rat aortic vascular smooth muscle cells via the activation of pro-inflammatory pathways (Mi et al. 2022) and was elevated in aged SAMP8 mouse muscle (Hoshino et al. 2022), suggesting both a mechanistic and biomarker role.

Galactosamine, identified as increased in our study, has been described as a metabolic inhibitor capable of inducing senescence in hepatocytes under chemically induced and oxidative stress conditions (Kumar et al. 2024), indicating that its accumulation may reflect impaired metabolic homeostasis in senescent cells. In addition, S-lactoylglutathione was elevated in our model, consistent with findings in replicative senescence of human fibroblasts, where its accumulation has been linked to increased methylglyoxal production and altered glycolytic flux (James et al. 2016).

Beyond lipid and amino acid metabolism, additional metabolites associated with senescence were identified. Uric acid levels were increased in our models of BEC senescence, consistent with reports in mouse lens epithelial cells, where uric acid accumulation promotes senescence and contributes to cataract formation via inflammasome activation (Lin et al. 2024). We also detected elevated levels of adenosine 5’monophosphate, in agreement with findings from skeletal muscle of aged (40- and 55- vs. 12-weeks-old) SAMP8 mice (Hoshino et al. 2022), further supporting conserved alterations in nucleotide metabolism across senescence models.

N-acetylneuraminate was increased in our dataset, linking our findings to sialic acid metabolism. Its derivative, N-acetylneuraminic acid (Neu5Ac), has been implicated in promoting inflammatory cytokine production in adipose tissue. Although not directly measured, genetic ablation of its downstream converting enzyme (cytidine monophosphate-N-acetylneuraminic acid hydroxylase) reduces senescence markers in mice aged 24-25 weeks (Takeda et al. 2023). This suggests that alterations in this pathway are functionally linked to senescence and may support a senescence-associated increase in upstream sialic acid metabolites.

Together, these findings indicate that while core metabolic pathways affected by senescence are conserved across models, the directionality of individual metabolites can vary depending on cell type, extracellular environment, and the specific senescence trigger.

### Identification of novel senescence-associated metabolites

In addition to metabolites already linked to senescence, our dataset revealed several novel candidates that were consistently altered by both inducers (threshold fold-change ≥ 2 for increased or ≤ 0.5 for decreased metabolites, **Table 2**). For instance, further support for altered fatty acid oxidation came from increased glutarylcarnitine and propionylcarnitine in the supernatant, while carnitine-18, a known senescence marker, was decreased in the cells. Additionally, ceramides (ceramide-27 and −29, and hexanoyl-2-ceramide-29) were decreased in cell lysates (**Fig. 3c**), supporting the idea that sphingolipid depletion is a hallmark of senescence and that ceramides are often considered inducers of senescence (Trayssac et al. 2018).

**Table 2.**
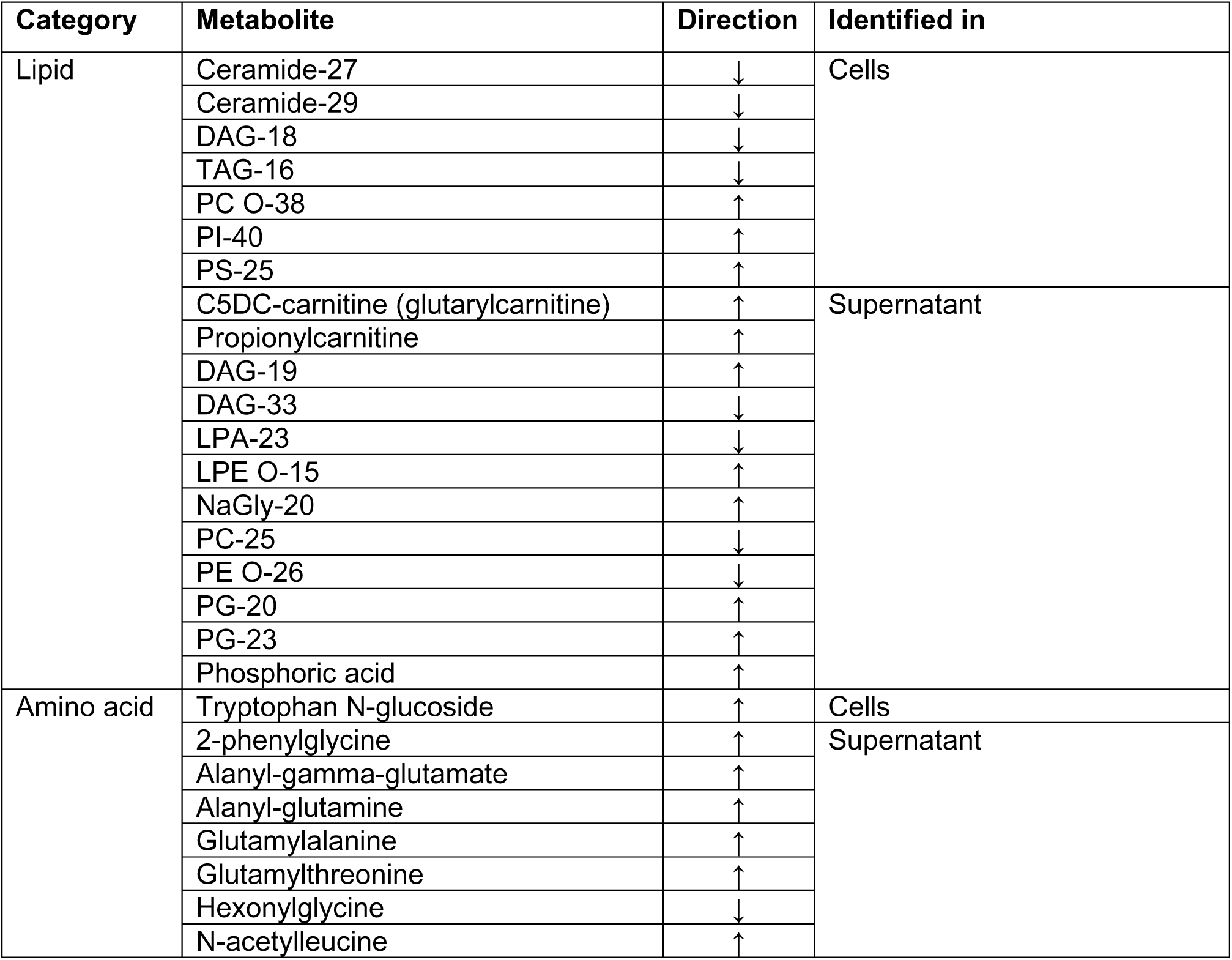
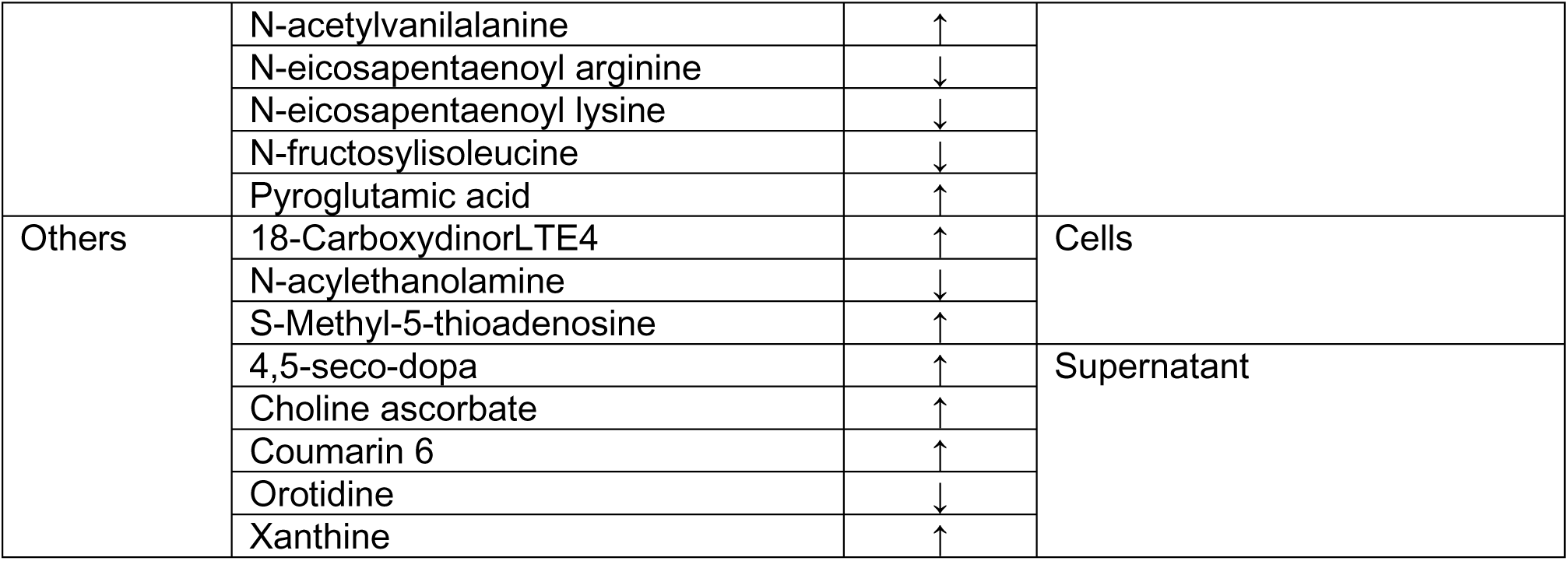
Novel potential senescence metabolites. Metabolites changed in the same direction in both Dox and H_2_O_2_ treatment with threshold fold-change ≥ 2 for increased or ≤ 0.5 for decreased metabolites. Directionality refers to changes in senescent versus control conditions. CAR = carnitine, DAG = diacylglycerides, LPA = lysophosphatidic acid, LPE = lysophosphatidylethanolamine, NAGly = N-acyl-glycine, PC = phosphatidylcholine, PE = phosphatidyl ethanolamine, PG = phosphaditylglycerol, PI = phosphatidylinositol, PS = phosphaditylserine, PC O = ether-linked phosphatidylcholine, PE O = ether-linked phosphatidylethanolamine, TAG = triacylglycerides.

We also observed distinct changes in glycerophospholipid species, beyond those previously reported. Elevated metabolites included PC O-38, PS-25, and PI-40 in cell lysates, while both increased (e.g., LPE O-15) and decreased (e.g., LPA-23, PC-25, PE O-26, PG-20, PG-23) glycerophospholipids were found in the supernatant. Additionally, DAGs – linked to cytokines of the senescence-associated secretory phenotype such as CCL2/MCP-1 (Hamsanathan et al. 2024) – showed divergent regulation: DAG-18 was decreased in cell lysates, while DAG-19 was increased and DAG-33 was decreased in the supernatant. This highlights the complexity of lipid remodeling in senescence. Interestingly, although TAG biosynthesis has been associated with chemotherapy-induced senescence of tumor cells via upregulation of DGAT1 (Hamsanathan and Gurkar 2022), we observed a decrease in TAG-16 in senescent BECs, suggesting cell-type-specific regulation.

Apart from lipid metabolites, several dipeptides and amino acid derivatives – including alanyl-gamma-glutamate, alanyl-glutamine, glutamylalanine, and glutamylthreonine – were found to be altered, potentially reflecting changes in protein turnover and amino acid recycling. These metabolites may arise from altered proteolysis or peptidase activity, which have been implicated in cellular stress responses that occur with aging including senescence (Frankowska et al. 2022).

Further supporting a shift in amino acid metabolism, we observed changes in N-acetylleucine, N-acetylvanilalanine, N-fructosylisoleucine, and tryptophan N-glucoside. Acetylation and glycosylation of amino acids are post-translational and metabolic modifications that can influence signaling, oxidative stress handling, and metabolic flux (Yang et al. 2023). While these metabolites have not been directly linked to senescence, their recurrent alteration across treatments points toward broader modulation of amino acid processing.

Notably, pyroglutamic acid was also among the altered metabolites, which may be linked to disrupted redox homeostasis, as its accumulation has been reported under conditions of glutathione depletion and oxidative stress (Gamarra et al. 2019) that are also hallmarks of senescence.

Changes were also observed in purine and pyrimidine metabolism, including increased levels of xanthine and S-methyl-5-thioadenosine whereas orotidine was decreased. S-methyl-5-thioadenosine is a byproduct of polyamine synthesis and has been shown to accumulate under conditions of oxidative stress or inflammation (Li et al. 2019). Decreased S-methyl-5-thioadenosine levels have been associated with impaired anti-inflammatory capacity and increased susceptibility to cognitive decline following surgery in elderly humans and aged mice (Zhang et al. 2024). Dysregulation of nucleotide metabolism is a known feature of senescent cells (Delfarah et al. 2019), potentially affecting DNA repair and epigenetic regulation.

Finally, the increase of 18-carboxydinorLTE4, a metabolite of leukotriene E4, suggests the engagement of inflammatory eicosanoid pathways. Elevated leukotriene derivatives have been linked to inflammatory signaling cascades commonly observed in diverse senescent cell types (Wiley et al. 2019; Wiley and Campisi 2021).

### Pathway mapping of senescent-related metabolites in BECs

To contextualize the identified metabolites within biological pathways, we qualitatively mapped them using MetaboAnalyst 6.0 and the Reactome Pathway Browser. This qualitative mapping highlighted three lipid-related pathways with the greatest coverage in our dataset (**Fig. 5**):

1) Fatty acid β-oxidation, indicated by changes in carnitine species. This process relies on membrane and mitochondrial transporters such as SLC22A5, which mediates carnitine uptake across the plasma membrane, and SLC25A20, which facilitates carnitine transport into mitochondria. Additionally, fatty acid transport proteins are involved in the cellular uptake of free fatty acids (Longo et al. 2016), which are subsequently converted to fatty acyl-CoA and then to acylcarnitine.
2) Sphingolipid metabolism, where sphingomyelins are hydrolyzed to form ceramides. Alternatively, ceramide synthesis via the *de novo* pathway involves several intermediates, including sphinganine (Obeid and Hannun 2003; SenthilKumar et al. 2024). These ceramides are known to play key roles in cell stress responses and are closely linked to senescence-associated signaling (Stith et al. 2019).
3) Glycerophospholipid metabolism – where the endoplasmic reticulum serves as the origin of many identified metabolites. In this pathway, glycerol-3-phosphate is the precursor for LPA, which is converted to phosphatidic acid by LPA acyltransferase. Phosphatidic acid is further transformed into DAG and subsequently PE, PC, and TAG, or alternatively into cytidine diphosphate-DAG, the precursor of PI, PG, cardiolipin, and bis(monoacylglycero)phosphate (Takeuchi and Reue 2009; Vance 2015).

**Figure 5:**
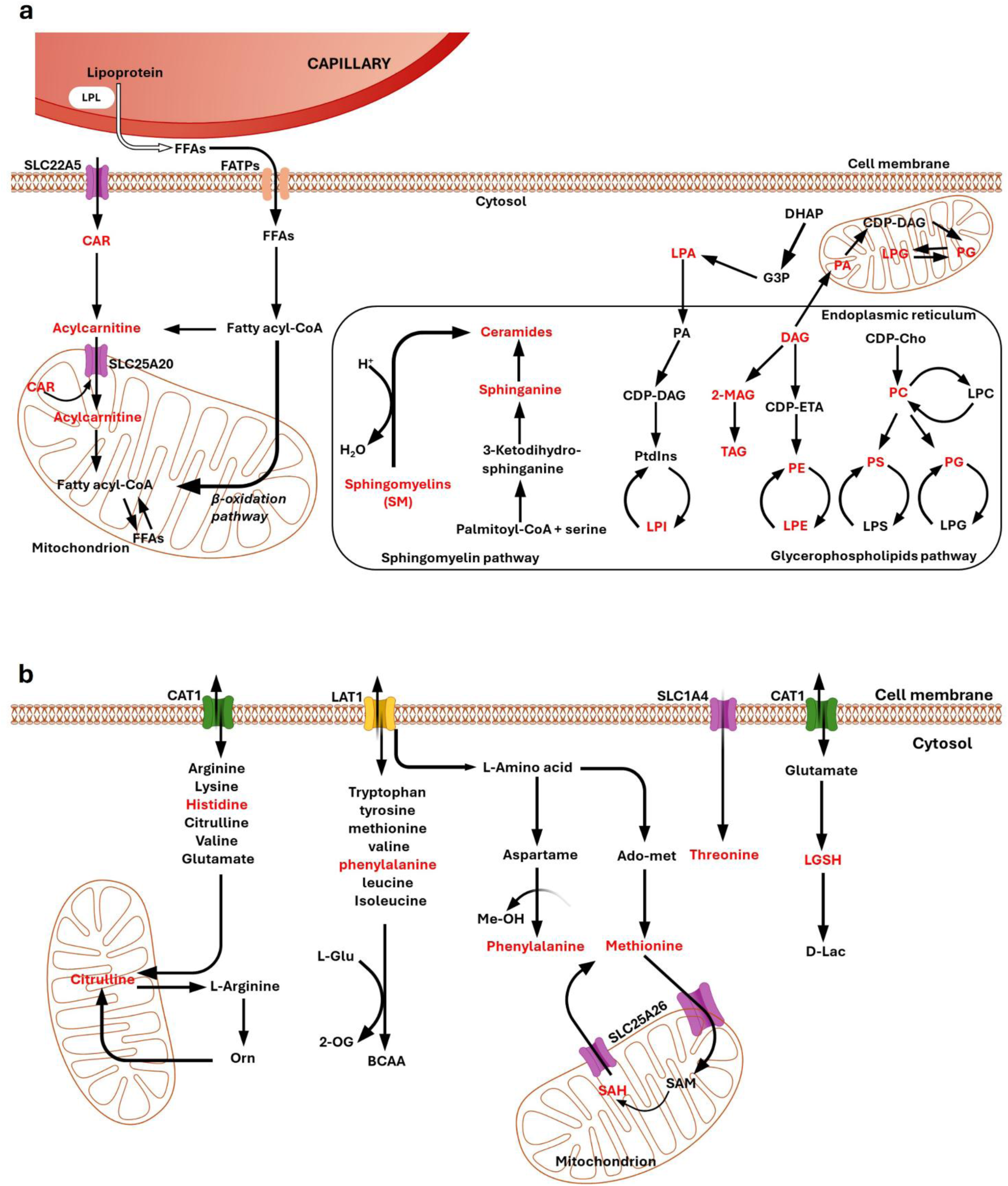
Identified senescence metabolites and their involvement in the metabolic pathways. **a)** Lipid metabolites involved in BEC senescence. CAR = carnitine, CDO-Cho = cytidine diphosphocholine, CDP-DAG = diphosphate-diacylglycerols, CDP-ETA = cytidine diphosphoethanolamine, DAG = diacylglyceride, DHAP = dihydroxy acetone phosphate, FFAs = free fatty acids, FATPs = fatty-acid transporter proteins, G3P = glycerol-3-phosphate, LPA = lysophosphatidic acid, LPC =lysophosphatidyl choline, LPE = lysophosphatidylethanolamine, LPG = lysophosphatidylglycerol, LPI = lysophosphatidylinositol, LPS = lysophosphatidyl serine, MAG = monoglyceride, PA = phosphatidic acid, PC = phosphatidylcholine, PE = phosphatidylethanolamine, PG = phosphatidylglycerol, PS = lysophosphatidylserine, PtdIns = phosphadityl inositol, TAG = triglyceride. **b)** Amino acid metabolites involved in BEC senescence. 2-OG = 2-oxoglutarate, Ado-met = adenosine methionine, BCAA = branched amino acid, CAT1 = cationic amino transporter, LAT1 = large amino acid transporter, L-Glu = L-glutamine, LGSH = lactoylglutathione, Me-OH = methanol, SAH = S-adenosylhomocysteine, SAM = S-adenosylmethionine, SLC = solute carrier, Val = valine. The red color indicates the specific metabolites identified in our pathway analysis.

Beyond lipid metabolism, we also identified significant alterations in amino acid metabolism. Notably, amino acid transport appeared altered, including:

1) cationic amino acid transporter 1 (CAT1), which primarily mediates the transport of arginine, lysine, histidine, citrulline, valine, and glutamate (Closs 2006; Jungnickel et al. 2018),
2) L-type amino acids transporter 1 (LAT1), responsible for the uptake of large neutral amino acids such as tryptophan, tyrosine, methionine, valine, phenylalanine, leucine and isoleucine (Yanagida et al. 2001), and
3) members of solute carrier (SLC) family, including SLC1A4, which is directly involved in the transport of threonine, among others.

These transport alterations reflect the broader reprogramming of amino acid availability during senescence. Interestingly, many of the detected amino acid metabolites in our dataset were acetylated forms, amino sugar conjugates, or small peptides. While these modified forms may not map directly onto canonical pathways, their precursors and processing intermediates align with known senescence-associated reprogramming.

In summary, these amino acid-related pathways, alongside lipid metabolism, reflect a broader metabolic shift associated with cellular senescence. Notably, several of the observed pathway alterations in our study are consistent with findings in other aging and senescence models (James et al. 2016; Hamsanathan et al. 2024), further supporting the relevance of our model to age-related metabolic dysregulation.

## Discussion

In this study, we successfully established *in vitro* models of BEC senescence using Dox or H_2_O_2_ as evidenced by increased SA-β-gal activity, p16, p21, and *Il-6* levels. We also document selective reductions in tight-junction proteins in these models. Using untargeted metabolomics and lipidomics, we present a novel dataset of the metabolites involved in BEC senescence. We were able to demonstrate common metabolites between both treatments that are likely related to BBB senescence and identify potential novel senescence metabolites.

Dox and H_2_O_2_ are widely recognized senescence inducers commonly used to investigate cellular senescence across various cell types (Kudlova et al. 2022). H_2_O_2_ treatment enhanced SA-β-gal activity, p16 and p21 expression in murine BECs, including cerebEND and cEND cell lines, as well as primary cells (Salvador et al. 2021). Similar effects have been observed in the human cerebro-microvascular endothelial cell line (hCMEC/D3) (Ting et al. 2023). Most research on Dox has focused on peripheral endothelial cells, demonstrating that it also raises SA-β-gal activity, p16 and p21 expression (Graziani et al. 2022; Abdelgawad et al. 2022). Therefore, our findings of elevated SA-β-gal activity, p16 and p21 expression in bEnd.3 cells following both treatments align with these previously published results.

Furthermore, the upregulation of the SASP cytokine *Il-6*, recapitulating prior findings in peripheral endothelial cells exposed to Dox (Abdelgawad et al. 2022), supports an inflammatory milieu that BECs can sense via cytokine receptors, promoting endothelial activation and context-dependent tight-junction remodeling (Blecharz-Lang et al. 2018; Acioglu and Elkabes 2025). In line with this, we observed a robust decrease in occludin, a significant H_2_O_2_-dependent decrease in ZO-1 with a similar trend under doxorubicin, and a variable, non-significant change in claudin-5. This pattern fits reports that physiological BBB aging often shows selective junctional changes and increased transcytosis rather than uniform tight junction loss (Knowland et al. 2014; Ben-Zvi et al. 2014; Yang et al. 2020; Cummins et al. 2024), with more consistent tight junction reductions under inflammatory stress. This pattern is consistent with prior *in vitro* senescence models in BECs, where H_2_O_2_ disrupted ZO-1 and reduced claudin-5 expression, although reductions in claudin-5 also did not reach significance in that study (Salvador et al. 2021). Our data therefore suggest that senescence-associated inflammatory signals (e.g., IL-6) may bias junctional remodeling in BECs, with occludin being particularly vulnerable under our conditions.

In our untargeted MS workflow, preliminary MS1-based identifications were performed with a strict mass accuracy threshold of ±5 ppm. We then curated the data by matching MS2 spectra to their most likely candidates using spectra mentioned in section **2.6.3 Identification.** Spectral alignment was carried out by comparing measured MS2 spectra to reference library entries, with each match evaluated using a dot product score ranging from 0 to 1, where 1 indicates a perfect spectral match.

It should be noted, however, that the high-resolution MS1- and MS2-based identification approach employed here has its limitations, particularly regarding the level of structural detail that can be confidently assigned. In several cases, especially within certain lipid subclasses such as DAGs, PCs and CARs; the available MS2 spectral information and library coverage did not allow for unambiguous identification at the isomeric or acyl chain double bond positional level. For example, features corresponding to PG 18:0 and PG 18:1 could not be reliably distinguished due to limited fragment ions. In such instances, we chose to report identifications at the lipid class and total carbon level (e.g. PG-18 as shown in the figures), acknowledging the presence of unresolved isobaric or isomeric species. While more advanced methodologies such as ion mobility spectrometry may offer higher structural resolution, these were beyond the scope and technical setup of the current study. Our approach, based on high-resolution mass spectrometry (HRMS) MS1 data and spectral matching against public libraries, was deemed sufficient for the biological objectives of this work, namely the generation of a consistent and interpretable molecular fingerprint of senescence-induced metabolic changes.

In conclusion, this study provides the first comprehensive metabolic characterization of senescence in BECs, a critical component of the BBB. Using two established senescence inducers, doxorubicin and hydrogen peroxide, we confirmed the induction of cellular senescence in mouse BECs through increased SA-β-gal activity and upregulated of p16 and p21 expression. Metabolomic profiling revealed distinct and overlapping changes in both intracellular and secreted metabolites, with several of these having been previously associated with aging and senescence in other tissues, while others representing novel candidates. Our findings shed light on the metabolic reprogramming that accompanies BEC senescence and provide valuable insights into mechanisms that may contribute to BBB dysfunction during aging. These results lay the groundwork for the development of metabolic biomarkers and potential therapeutic targets to mitigate BBB aging and its contribution to neurodegenerative diseases such as Alzheimer’s disease and vascular dementia.

## Limitations and future directions

Several considerations limit the interpretation and generalizability of our findings. First, the use of an immortalized mouse BEC line in monoculture does not recapitulate neurovascular unit interactions, basement membrane context, or hemodynamic forces present *in vivo*. These elements are known to modulate tight-junction organization, vesicular transport, and glial-endothelial signaling and are better captured in multicellular, flow-based BBB models. We therefore provided a staged plan for validation in multicellular/flow-based BBB models and *in vivo*. Second, senescence was induced using two stressors (doxorubicin and H_2_O_2_), which strengthens inference about shared signatures but does not encompass the full spectrum of senescence programs across brain regions or species. Third, although we provide protein-level evidence of tight-junction alterations, we did not assess efflux transporter function in this study, and *in vivo* validation across ages was beyond scope. Finally, untargeted MS-based identification has limited isomeric resolution for some lipid subclasses, which we explicitly report and curate at the lipid-class/total-carbon level when needed.

Within this context, a staged translational roadmap can help bridge from discovery to *in vivo* relevance. Prioritized metabolite candidates and pathways could be evaluated in advanced multicellular BBB models (e.g., co/tri-cultures or humanized, flow-based platforms) that more closely reproduce neurovascular unit architecture and shear stress (Salman et al. 2020; O’Halloran et al. 2025). Mechanism-focused perturbations (e.g., metabolite add-back/inhibition, receptor or transporter modulation) under oxidative or inflammatory stress would further test pathway necessity and sufficiency in a more physiologically relevant context. Subsequent *in vivo* studies in aged mice could then deploy complementary BBB readouts that index distinct barrier mechanisms, recognizing that different measures (e.g., small-molecule tracers versus protein leakage) capture paracellular versus transcellular changes and may not fully overlap (Cummins et al. 2024). This stepwise path, from monoculture discovery to multicellular/flow BBB validation and on to *in vivo* confirmation, is intended to enhance mechanistic resolution, reproducibility, and causal inference prior to human translation.

## Appendix. Supplementary Material

Figures S1-S4, Supplementary Tables 1-2, see attached

## Data Availability

All data have been deposited in the Center for Computational Mass Spectrometry’s MassIVE data repository. The DOI will be available upon acceptance of the manuscript.

## List of abbreviations

BBB: Blood-brain barrier
BECs: Brain endothelial cells
CAR: Carnitine
DAG: Diacylglyceride
Dox: Doxorubicin
H_2_O_2_: Hydrogen peroxide
HRMS: High-resolution mass spectrometry
LPE: Lysophosphatidylethanolamine
LPI: Lysophosphatidylinositol
MS: Mass spectrometry
PLSDA: Partial least squares discriminant analysis
PCR: Polymerase chain reaction
SA-β-gal: Senescence-associated β-galactosidase
SASP: Senescence-associated secretory phenotype
TAG: Triacylglyceride
ZO-1: Zonula occludens 1

## Authors’ contributions

**Ammar Tahir:** Writing – original draft, review & editing, Conceptualization, Data curation, Metabolomics methodology and analysis, Resources. **Hari Baskar Balasubramanian:** Writing – original draft, review & editing, Investigation, Formal analysis, Visualization. **Dominik Kahr:** Writing – review & editing, Investigation, Formal analysis. **Florian Haage:** Writing – review & editing, Investigation, Formal analysis. **Marietta Zille:** Writing – original draft, review & editing, Supervision, Formal analysis, Conceptualization, Resources, Project administration.

## Acknowledgements

The authors thank Andrea Szabo for the professional support in creating and improving Figures 5 and S1.

## References

Abdelgawad IY, Agostinucci K, Ismail SG, Grant MKO, Zordoky BN (2022) EA.hy926 Cells and HUVECs Share Similar Senescence Phenotypes but Respond Differently to the Senolytic Drug ABT-263. Cells 11(13):1992. 10.3390/cells11131992

Acioglu C, Elkabes S (2025) Innate immune sensors and regulators at the blood brain barrier: focus on toll-like receptors and inflammasomes as mediators of neuro-immune crosstalk and inflammation. J Neuroinflammation 22(1):39. 10.1186/s12974-025-03360-3

Archie SR, Al Shoyaib A, Cucullo L (2021) Blood-Brain Barrier Dysfunction in CNS Disorders and Putative Therapeutic Targets: An Overview. Pharmaceutics 13(11):1779. 10.3390/pharmaceutics13111779

Ballabh P, Braun A, Nedergaard M (2004) The blood–brain barrier: an overview. Neurobiol Dis 16(1):1–13. 10.1016/j.nbd.2003.12.016

Banks WA, Reed MJ, Logsdon AF, Rhea EM, Erickson MA (2021) Healthy aging and the blood–brain barrier. Nat Aging 1(3):243–254. 10.1038/s43587-021-00043-5

Ben-Zvi A, Lacoste B, Kur E, Andreone BJ, Mayshar Y, Yan H, Gu C (2014) Mfsd2a is critical for the formation and function of the blood–brain barrier. Nature 509(7501):507–511. 10.1038/nature13324

Bhalerao A, Sivandzade F, Archie SR, Chowdhury EA, Noorani B, Cucullo L (2020) In vitro modeling of the neurovascular unit: advances in the field. Fluids Barriers CNS 17(1):22. 10.1186/s12987-020-00183-7

Blecharz-Lang KG, Wagner J, Fries A, Nieminen-Kelhä M, Rösner J, Schneider UC, Vajkoczy P (2018) Interleukin 6-Mediated Endothelial Barrier Disturbances Can Be Attenuated by Blockade of the IL6 Receptor Expressed in Brain Microvascular Endothelial Cells. Transl Stroke Res 9(6):631–642. 10.1007/s12975-018-0614-2

Cadenas C, Vosbeck S, Hein E-M, Hellwig B, Langer A, Hayen H, Franckenstein D, Büttner B, Hammad S, Marchan R, Hermes M, Selinski S, Rahnenführer J, Peksel B, Török Z, Vígh L, Hengstler JG (2012) Glycerophospholipid profile in oncogene-induced senescence. Biochim Biophys Acta BBA - Mol Cell Biol Lipids 1821(9):1256–1268. 10.1016/j.bbalip.2011.11.008

Campisi J (2005) Senescent Cells, Tumor Suppression, and Organismal Aging: Good Citizens, Bad Neighbors. Cell 120(4):513–522. 10.1016/j.cell.2005.02.003

Castro A, Signini ÉF, De Oliveira JM, Di Medeiros Leal MCB, Rehder-Santos P, Millan-Mattos JC, Minatel V, Pantoni CBF, Oliveira RV, Catai AM, Ferreira AG (2022) The Aging Process: A Metabolomics Perspective. Molecules 27(24):8656. 10.3390/molecules27248656

Chaulagain B, Gothwal A, Lamptey RNL, Trivedi R, Mahanta AK, Layek B, Singh J (2023) Experimental Models of In Vitro Blood–Brain Barrier for CNS Drug Delivery: An Evolutionary Perspective. Int J Mol Sci 24(3):2710. 10.3390/ijms24032710

Closs EI (2006) Structure and Function of Cationic Amino Acid Transporters (CATs). 213(2):67–77. doi:10.1007/s00232-006-0875-7

Cummins MJ, Cresswell ET, Bevege RJ, Smith DW (2024) Aging disrupts blood–brain and blood-spinal cord barrier homeostasis, but does not increase paracellular permeability. GeroScience 47(1):263–285. 10.1007/s11357-024-01404-9

Debacq-Chainiaux F, Erusalimsky JD, Campisi J, Toussaint O (2009) Protocols to detect senescence-associated beta-galactosidase (SA-βgal) activity, a biomarker of senescent cells in culture and in vivo. Nat Protoc 4(12):1798–1806. 10.1038/nprot.2009.191

Delfarah A, Parrish S, Junge JA, Yang J, Seo F, Li S, Mac J, Wang P, Fraser SE, Graham NA (2019) Inhibition of nucleotide synthesis promotes replicative senescence of human mammary epithelial cells. J Biol Chem 294(27):10564–10578. 10.1074/jbc.RA118.005806

Erdő F, Denes L, De Lange E (2017) Age-associated physiological and pathological changes at the blood–brain barrier: A review. J Cereb Blood Flow Metab 37(1):4–24. 10.1177/0271678X16679420

Fang W, Chen S, Jin X, Liu S, Cao X, Liu B (2023) Metabolomics in aging research: aging markers from organs. Front Cell Dev Biol 11:1198794. 10.3389/fcell.2023.1198794

Frankowska N, Lisowska K, Witkowski JM (2022) Proteolysis dysfunction in the process of aging and age-related diseases. Front Aging 3:927630. 10.3389/fragi.2022.927630

Gamarra Y, Santiago FC, Molina-López J, Castaño J, Herrera-Quintana L, Domínguez Á, Planells E (2019) Pyroglutamic acidosis by glutathione regeneration blockage in critical patients with septic shock. Crit Care 23(1):162. 10.1186/s13054-019-2450-5

Giuliani A, Giudetti AM, Vergara D, Del Coco L, Ramini D, Caccese S, Sbriscia M, Graciotti L, Fulgenzi G, Tiano L, Fanizzi FP, Olivieri F, Rippo MR, Sabbatinelli J (2023) Senescent Endothelial Cells Sustain Their Senescence-Associated Secretory Phenotype (SASP) through Enhanced Fatty Acid Oxidation. Antioxidants 12(11):1956. 10.3390/antiox12111956

Graziani S, Scorrano L, Pontarin G (2022) Transient Exposure of Endothelial Cells to Doxorubicin Leads to Long-Lasting Vascular Endothelial Growth Factor Receptor 2 Downregulation. Cells 11(2):210. 10.3390/cells11020210

Hamsanathan S, Anthonymuthu T, Prosser D, Lokshin A, Greenspan SL, Resnick NM, Perera S, Okawa S, Narasimhan G, Gurkar AU (2024) A molecular index for biological age identified from the metabolome and senescence-associated secretome in humans. Aging Cell 23(4):e14104. 10.1111/acel.14104

Hamsanathan S, Gurkar AU (2022) Lipids as Regulators of Cellular Senescence. Front Physiol 13:796850. 10.3389/fphys.2022.796850

Han X, Gross RW (2022) The foundations and development of lipidomics. J Lipid Res 63(2):100164. 10.1016/j.jlr.2021.100164

Helman A, Klochendler A, Azazmeh N, Gabai Y, Horwitz E, Anzi S, Swisa A, Condiotti R, Granit RZ, Nevo Y, Fixler Y, Shreibman D, Zamir A, Tornovsky-Babeay S, Dai C, Glaser B, Powers AC, Shapiro AMJ, Magnuson MA, Dor Y, Ben-Porath I (2016) p16Ink4a-induced senescence of pancreatic beta cells enhances insulin secretion. Nat Med 22(4):412–420. 10.1038/nm.4054

Hoshino T, Kato Y, Sugahara K, Katakura A (2022) Aging-related metabolic changes in the extensor digitorum longus muscle of senescence-accelerated mouse-prone 8. Geriatr Gerontol Int 22(2):160–167. 10.1111/ggi.14333

James EL, Lane JAE, Michalek RD, Karoly ED, Parkinson EK (2016) Replicatively senescent human fibroblasts reveal a distinct intracellular metabolic profile with alterations in NAD+ and nicotinamide metabolism. Sci Rep 6(1):38489. 10.1038/srep38489

Johnson CH, Ivanisevic J, Siuzdak G (2016) Metabolomics: beyond biomarkers and towards mechanisms. Nat Rev Mol Cell Biol 17(7):451–459. 10.1038/nrm.2016.25

Jungnickel KEJ, Parker JL, Newstead S (2018) Structural basis for amino acid transport by the CAT family of SLC7 transporters. Nat Commun 9(1):550. 10.1038/s41467-018-03066-6

Kind T, Liu K-H, Lee DY, DeFelice B, Meissen JK, Fiehn O (2013) LipidBlast in silico tandem mass spectrometry database for lipid identification. Nat Methods 10(8):755–758. 10.1038/nmeth.2551

Kiss T, Nyúl-Tóth Á, Balasubramanian P, Tarantini S, Ahire C, DelFavero J, Yabluchanskiy A, Csipo T, Farkas E, Wiley G, Garman L, Csiszar A, Ungvari Z (2020) Single-cell RNA sequencing identifies senescent cerebromicrovascular endothelial cells in the aged mouse brain. GeroScience 42(2):429–444. 10.1007/s11357-020-00177-1

Knopp RC, Erickson MA, Rhea EM, Reed MJ, Banks WA (2023) Cellular senescence and the blood–brain barrier: Implications for aging and age-related diseases. Exp Biol Med 248(5):399–411. 10.1177/15353702231157917

Knowland D, Arac A, Sekiguchi KJ, Hsu M, Lutz SE, Perrino J, Steinberg GK, Barres BA, Nimmerjahn A, Agalliu D (2014) Stepwise Recruitment of Transcellular and Paracellular Pathways Underlies Blood-Brain Barrier Breakdown in Stroke. Neuron 82(3):603–617. 10.1016/j.neuron.2014.03.003

Knox EG, Aburto MR, Clarke G, Cryan JF, O’Driscoll CM (2022) The blood-brain barrier in aging and neurodegeneration. Mol Psychiatry 27(6):2659–2673. 10.1038/s41380-022-01511-z

Kudlova N, De Sanctis JB, Hajduch M (2022) Cellular Senescence: Molecular Targets, Biomarkers, and Senolytic Drugs. Int J Mol Sci 23(8):4168. 10.3390/ijms23084168

Kumar P, Hassan M, Tacke F, Engelmann C (2024) Delineating the heterogeneity of senescence-induced-functional alterations in hepatocytes. Cell Mol Life Sci 81(1):200. 10.1007/s00018-024-05230-2

Lai Z, Tsugawa H, Wohlgemuth G, Mehta S, Mueller M, Zheng Y, Ogiwara A, Meissen J, Showalter M, Takeuchi K, Kind T, Beal P, Arita M, Fiehn O (2018) Identifying metabolites by integrating metabolome databases with mass spectrometry cheminformatics. Nat Methods 15(1):53–56. 10.1038/nmeth.4512

Li Y, Wang Y, Wu P (2019) 5’-Methylthioadenosine and Cancer: old molecules, new understanding. J Cancer 10(4):927–936. 10.7150/jca.27160

Lin HL, Wang S, Sato K, Zhang YQ, He BT, Xu J, Nakazawa T, Qin YJ, Zhang HY (2024) Uric acid–driven NLRP3 inflammasome activation triggers lens epithelial cell senescence and cataract formation. Cell Death Discov 10(1):126. 10.1038/s41420-024-01900-z

Longo N, Frigeni M, Pasquali M (2016) Carnitine transport and fatty acid oxidation. Biochim Biophys Acta BBA - Mol Cell Res 1863(10):2422–2435. 10.1016/j.bbamcr.2016.01.023

López-Otín C, Blasco MA, Partridge L, Serrano M, Kroemer G (2023) Hallmarks of aging: An expanding universe. Cell 186(2):243–278. 10.1016/j.cell.2022.11.001

Marques L, Johnson AA, Stolzing A (2020) Doxorubicin generates senescent microglia that exhibit altered proteomes, higher levels of cytokine secretion, and a decreased ability to internalize amyloid β. Exp Cell Res 395(2):112203. 10.1016/j.yexcr.2020.112203

Mi J, Chen X, Yiran Y, Tang Y, Liu Q, Xiao J, Ling W (2022) S-adenosylhomocysteine induces cellular senescence in rat aorta vascular smooth muscle cells via NF-κB-SASP pathway. J Nutr Biochem 107:109063. 10.1016/j.jnutbio.2022.109063

Milacic M, Beavers D, Conley P, Gong C, Gillespie M, Griss J, Haw R, Jassal B, Matthews L, May B, Petryszak R, Ragueneau E, Rothfels K, Sevilla C, Shamovsky V, Stephan R, Tiwari K, Varusai T, Weiser J, Wright A, Wu G, Stein L, Hermjakob H, D’Eustachio P (2024) The Reactome Pathway Knowledgebase 2024. Nucleic Acids Res 52(D1):D672–D678. 10.1093/nar/gkad1025

Muñoz-Espín D, Serrano M (2014) Cellular senescence: from physiology to pathology. Nat Rev Mol Cell Biol 15(7):482–496. 10.1038/nrm3823

Novo JP, Gee L, Caetano CA, Tomé I, Vilaça A, Von Zglinicki T, Moreira IS, Jurk D, Rosa S, Ferreira L (2024) Blood–brain barrier dysfunction in aging is mediated by brain endothelial senescence. Aging Cell 23(9):e14270. 10.1111/acel.14270

Obeid LM, Hannun YA (2003) Ceramide, Stress, and a “LAG” in Aging. Sci Aging Knowl Environ 2003(39). 10.1126/sageke.2003.39.pe27

O’Halloran L, Akinsete O, Kogan AL, Wrona M, Mahdi AF (2025) 3D in vitro blood–brain barrier models: recent advances and their role in brain disease research and therapy. Front Pharmacol 16:1637602. 10.3389/fphar.2025.1637602

Pang Z, Chong J, Zhou G, de Lima Morais DA, Chang L, Barrette M, Gauthier C, Jacques P-É, Li S, Xia J (2021) MetaboAnalyst 5.0: narrowing the gap between raw spectra and functional insights. Nucleic Acids Res 49(W1):W388–W396. 10.1093/nar/gkab382

Panyard DJ, Yu B, Snyder MP (2022) The metabolomics of human aging: Advances, challenges, and opportunities. Sci Adv 8(42):eadd6155. 10.1126/sciadv.add6155

Real MGC, Falcione SR, Boghozian R, Clarke M, Todoran R, St Pierre A, Zhang Y, Joy T, Jickling GC (2024) Endothelial Cell Senescence Effect on the Blood-Brain Barrier in Stroke and Cognitive Impairment. Neurology 103(11):e210063. 10.1212/WNL.0000000000210063

Roninson IB (2003) Tumor Cell Senescence in Cancer Treatment. Cancer Res 63(11):2705–2715

Sabbatinelli J, Prattichizzo F, Olivieri F, Procopio AD, Rippo MR, Giuliani A (2019) Where Metabolism Meets Senescence: Focus on Endothelial Cells. Front Physiol 10:1523. 10.3389/fphys.2019.01523

Salman MM, Marsh G, Kusters I, Delincé M, Di Caprio G, Upadhyayula S, De Nola G, Hunt R, Ohashi KG, Gray T, Shimizu F, Sano Y, Kanda T, Obermeier B, Kirchhausen T (2020) Design and Validation of a Human Brain Endothelial Microvessel-on-a-Chip Open Microfluidic Model Enabling Advanced Optical Imaging. Front Bioeng Biotechnol 8:573775. 10.3389/fbioe.2020.573775

Salvador E, Burek M, Löhr M, Nagai M, Hagemann C, Förster CY (2021) Senescence and associated blood–brain barrier alterations in vitro. Histochem Cell Biol 156(3):283–292. 10.1007/s00418-021-01992-z

SenthilKumar G, Zirgibel Z, Cohen KE, Katunaric B, Jobe AM, Shult CG, Limpert RH, Freed JK (2024) Ying and Yang of Ceramide in the Vascular Endothelium. 44(8):1725–1736. doi:10.1161/ATVBAHA.124.321158

Sharma R, Diwan B (2023) Lipids and the hallmarks of ageing: From pathology to interventions. Mech Ageing Dev 215:111858. 10.1016/j.mad.2023.111858

Stith JL, Velazquez FN, Obeid LM (2019) Advances in determining signaling mechanisms of ceramide and role in disease. J Lipid Res 60(5):913–918. 10.1194/jlr.S092874

Tahir A, Draxler A, Stelzer T, Blaschke A, Laky B, Széll M, Binar J, Bartak V, Bragagna L, Maqboul L, Herzog T, Thell R, Wagner K-H (2024) A comprehensive IDA and SWATH-DIA Lipidomics and Metabolomics dataset: SARS-CoV-2 case control study. Sci Data 11(1):998. 10.1038/s41597-024-03822-y

Takeda R, Tabuchi A, Nonaka Y, Kano R, Sudo M, Kano Y, Hoshino D (2023) Cmah deficiency blunts cellular senescence in adipose tissues and improves whole-body glucose metabolism in aged mice. Geriatr Gerontol Int 23(12):958–964. 10.1111/ggi.14732

Takeuchi K, Reue K (2009) Biochemistry, physiology, and genetics of GPAT, AGPAT, and lipin enzymes in triglyceride synthesis. Am J Physiol-Endocrinol Metab 296(6):E1195–E1209. 10.1152/ajpendo.90958.2008

Ting KK, Coleman P, Kim HJ, Zhao Y, Mulangala J, Cheng NC, Li W, Gunatilake D, Johnstone DM, Loo L, Neely GG, Yang P, Götz J, Vadas MA, Gamble JR (2023) Vascular senescence and leak are features of the early breakdown of the blood–brain barrier in Alzheimer’s disease models. GeroScience 45(6):3307–3331. 10.1007/s11357-023-00927-x

Trayssac M, Hannun YA, Obeid LM (2018) Role of sphingolipids in senescence: implication in aging and age-related diseases. J Clin Invest 128(7):2702–2712. 10.1172/JCI97949

Tsugawa H, Ikeda K, Takahashi M, Satoh A, Mori Y, Uchino H, Okahashi N, Yamada Y, Tada I, Bonini P, Higashi Y, Okazaki Y, Zhou Z, Zhu Z-J, Koelmel J, Cajka T, Fiehn O, Saito K, Arita M, Arita M (2020) A lipidome atlas in MS-DIAL 4. Nat Biotechnol 38(10):1159–1163. 10.1038/s41587-020-0531-2

Vance JE (2015) Phospholipid Synthesis and Transport in Mammalian Cells. 16(1):1–18. doi:10.1111/tra.12230

Wang R, Li B, Lam SM, Shui G (2020) Integration of lipidomics and metabolomics for in-depth understanding of cellular mechanism and disease progression. J Genet Genomics 47(2):69–83. 10.1016/j.jgg.2019.11.009

Wiley CD, Brumwell AN, Davis SS, Jackson JR, Valdovinos A, Calhoun C, Alimirah F, Castellanos CA, Ruan R, Wei Y, Chapman HA, Ramanathan A, Campisi J, Jourdan Le Saux C (2019) Secretion of leukotrienes by senescent lung fibroblasts promotes pulmonary fibrosis. JCI Insight 4(24):e130056. 10.1172/jci.insight.130056

Wiley CD, Campisi J (2021) The metabolic roots of senescence: mechanisms and opportunities for intervention. Nat Metab 3(10):1290–1301. 10.1038/s42255-021-00483-8

Wishart DS, Tzur D, Knox C, Eisner R, Guo AC, Young N, Cheng D, Jewell K, Arndt D, Sawhney S, Fung C, Nikolai L, Lewis M, Coutouly M-A, Forsythe I, Tang P, Shrivastava S, Jeroncic K, Stothard P, Amegbey G, Block D, Hau DavidD, Wagner J, Miniaci J, Clements M, Gebremedhin M, Guo N, Zhang Y, Duggan GE, MacInnis GD, Weljie AM, Dowlatabadi R, Bamforth F, Clive D, Greiner R, Li L, Marrie T, Sykes BD, Vogel HJ, Querengesser L (2007) HMDB: the Human Metabolome Database. Nucleic Acids Res 35(Database):D521–D526. 10.1093/nar/gkl923

Xia J, Wishart DS (2010a) MSEA: a web-based tool to identify biologically meaningful patterns in quantitative metabolomic data. Nucleic Acids Res 38(Web Server):W71–W77. 10.1093/nar/gkq329

Xia J, Wishart DS (2010b) MetPA: a web-based metabolomics tool for pathway analysis and visualization. Bioinformatics 26(18):2342–2344. 10.1093/bioinformatics/btq418

Yanagida O, Kanai Y, Chairoungdua A, Kim DK, Segawa H, Nii T, Cha SH, Matsuo H, Fukushima J, Fukasawa Y, Tani Y, Taketani Y, Uchino H, Kim JY, Inatomi J, Okayasu I, Miyamoto K, Takeda E, Goya T, Endou H (2001) Human L-type amino acid transporter 1 (LAT1): characterization of function and expression in tumor cell lines. Biochim Biophys Acta BBA - Biomembr 1514(2):291–302. 10.1016/S0005-2736(01)00384-4

Yang AC, Stevens MY, Chen MB, Lee DP, Stähli D, Gate D, Contrepois K, Chen W, Iram T, Zhang L, Vest RT, Chaney A, Lehallier B, Olsson N, Du Bois H, Hsieh R, Cropper HC, Berdnik D, Li L, Wang EY, Traber GM, Bertozzi CR, Luo J, Snyder MP, Elias JE, Quake SR, James ML, Wyss-Coray T (2020) Physiological blood–brain transport is impaired with age by a shift in transcytosis. Nature 583(7816):425–430. 10.1038/s41586-020-2453-z

Yang Y-H, Wen R, Yang N, Zhang T-N, Liu C-F (2023) Roles of protein post-translational modifications in glucose and lipid metabolism: mechanisms and perspectives. Mol Med 29(1):93. 10.1186/s10020-023-00684-9

Yao Z, Van Velthoven CTJ, Kunst M, Zhang M, McMillen D, Lee C, Jung W, Goldy J, Abdelhak A, Aitken M, Baker K, Baker P, Barkan E, Bertagnolli D, Bhandiwad A, Bielstein C, Bishwakarma P, Campos J, Carey D, Casper T, Chakka AB, Chakrabarty R, Chavan S, Chen M, Clark M, Close J, Crichton K, Daniel S, DiValentin P, Dolbeare T, Ellingwood L, Fiabane E, Fliss T, Gee J, Gerstenberger J, Glandon A, Gloe J, Gould J, Gray J, Guilford N, Guzman J, Hirschstein D, Ho W, Hooper M, Huang M, Hupp M, Jin K, Kroll M, Lathia K, Leon A, Li S, Long B, Madigan Z, Malloy J, Malone J, Maltzer Z, Martin N, McCue R, McGinty R, Mei N, Melchor J, Meyerdierks E, Mollenkopf T, Moonsman S, Nguyen TN, Otto S, Pham T, Rimorin C, Ruiz A, Sanchez R, Sawyer L, Shapovalova N, Shepard N, Slaughterbeck C, Sulc J, Tieu M, Torkelson A, Tung H, Valera Cuevas N, Vance S, Wadhwani K, Ward K, Levi B, Farrell C, Young R, Staats B, Wang M-QM, Thompson CL, Mufti S, Pagan CM, Kruse L, Dee N, Sunkin SM, Esposito L, Hawrylycz MJ, Waters J, Ng L, Smith K, Tasic B, Zhuang X, Zeng H (2023) A high-resolution transcriptomic and spatial atlas of cell types in the whole mouse brain. Nature 624(7991):317–332. 10.1038/s41586-023-06812-z

Zhang L, Mao H, Zhou R, Zhu J, Wang H, Miao Z, Chen X, Yan J, Jiang H (2024) Low blood S-methyl-5-thioadenosine is associated with postoperative delayed neurocognitive recovery. Commun Biol 7(1):1356. 10.1038/s42003-024-07086-5

