## Appendix for "Aging-associated metabolomic and lipidomic remodeling in mouse brain endothelial cell senescence *in vitro*"

[†] Authors contributed equally

[*] Corresponding authors

Contact:

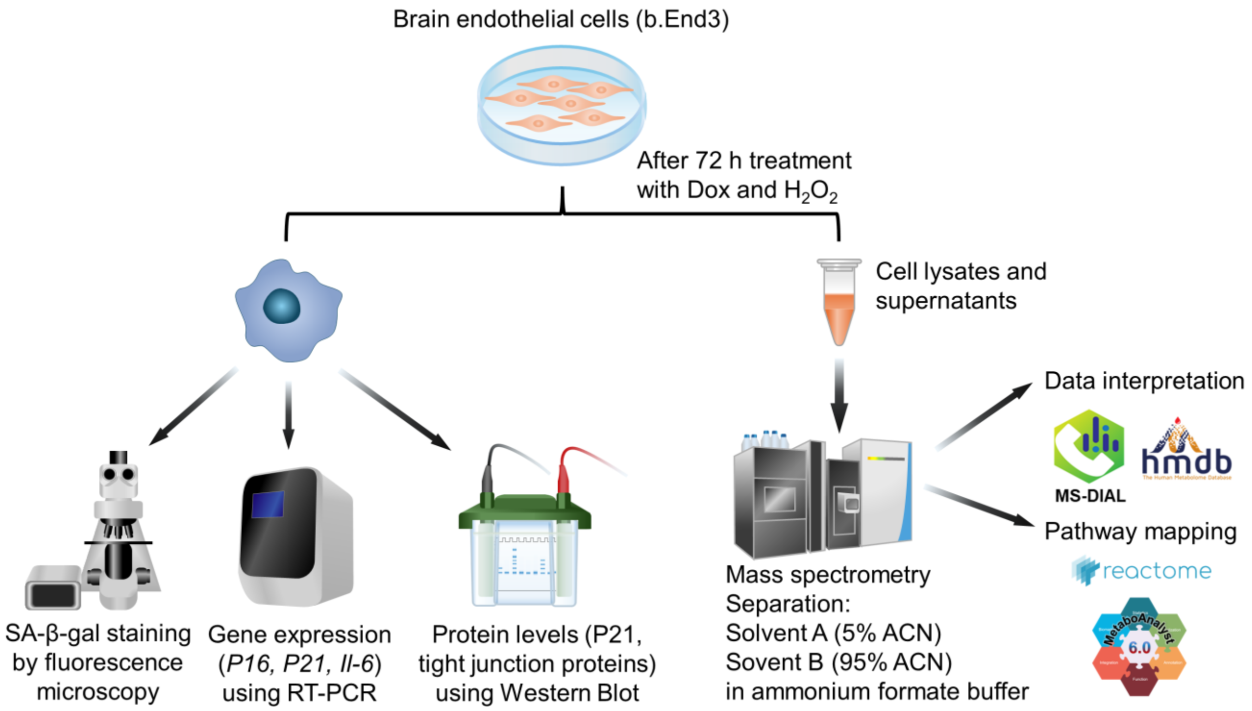


**Figure S1: Summary of the experimental design of the study.** Senescence was induced in bEnd.3 cells by 72-hours incubation with either doxorubicin or $H_2O_2$. We performed SA-β-gal imaging using fluorescence microscopy, RT-PCR to detect the relative gene expression of *P16*, *P21*, and *Il-6*, and Western Blotting to quantify p21 as well as the tight junction proteins ZO-1, occludin, and claudin-5. Cell lysates and supernatants were collected and analyzed using mass spectrometry. MS-DIAL, HMDB, and MetaboAnalyst were used to identify the metabolites and Reactome was used to perform pathway mapping.

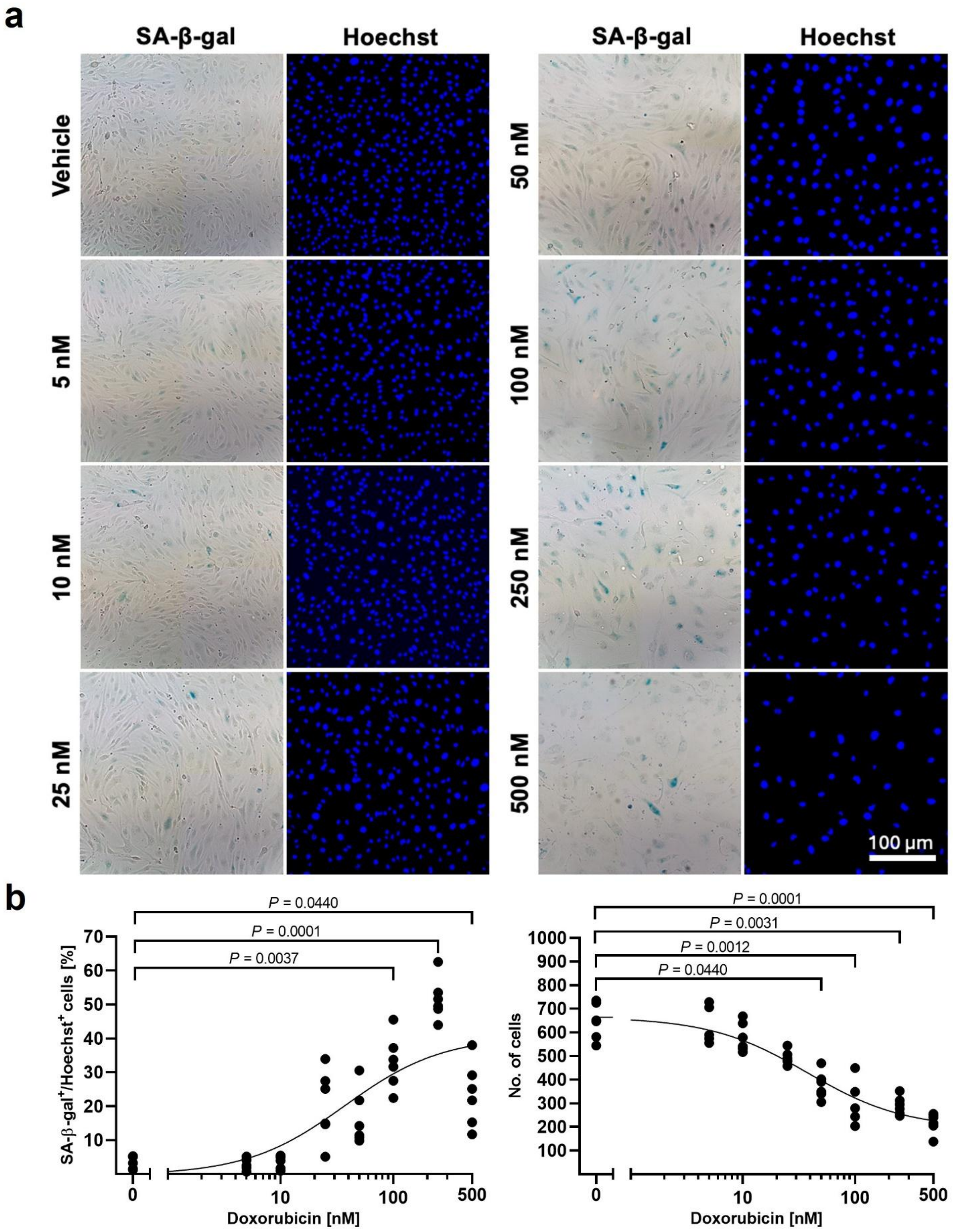


**Figure S2: Concentration-dependent increase in SA-β-gal activity after Dox treatment. A.** The SA-β-gal staining was imaged using bright-field microscopy. Fluorescence nuclear staining with Hoechst was performed to determine the total number of cells. **B.** The left graph shows the proportion of SA-β-gal positive cells at 72 hours of treatment. Kruskal-Wallis test: $\chi^2(8, N = 48) = 39.77$, $P < 0.0001$, $\eta^2 = 0.846$, followed by Dunn's post hoc test. The right graph represents the change in the total number of cells per concentration. Kruskal-Wallis test: $\chi^2(8, N = 48) = 41.84$, $P < 0.0001$, $\eta^2 = 0.890$, followed by Dunn's post hoc test.

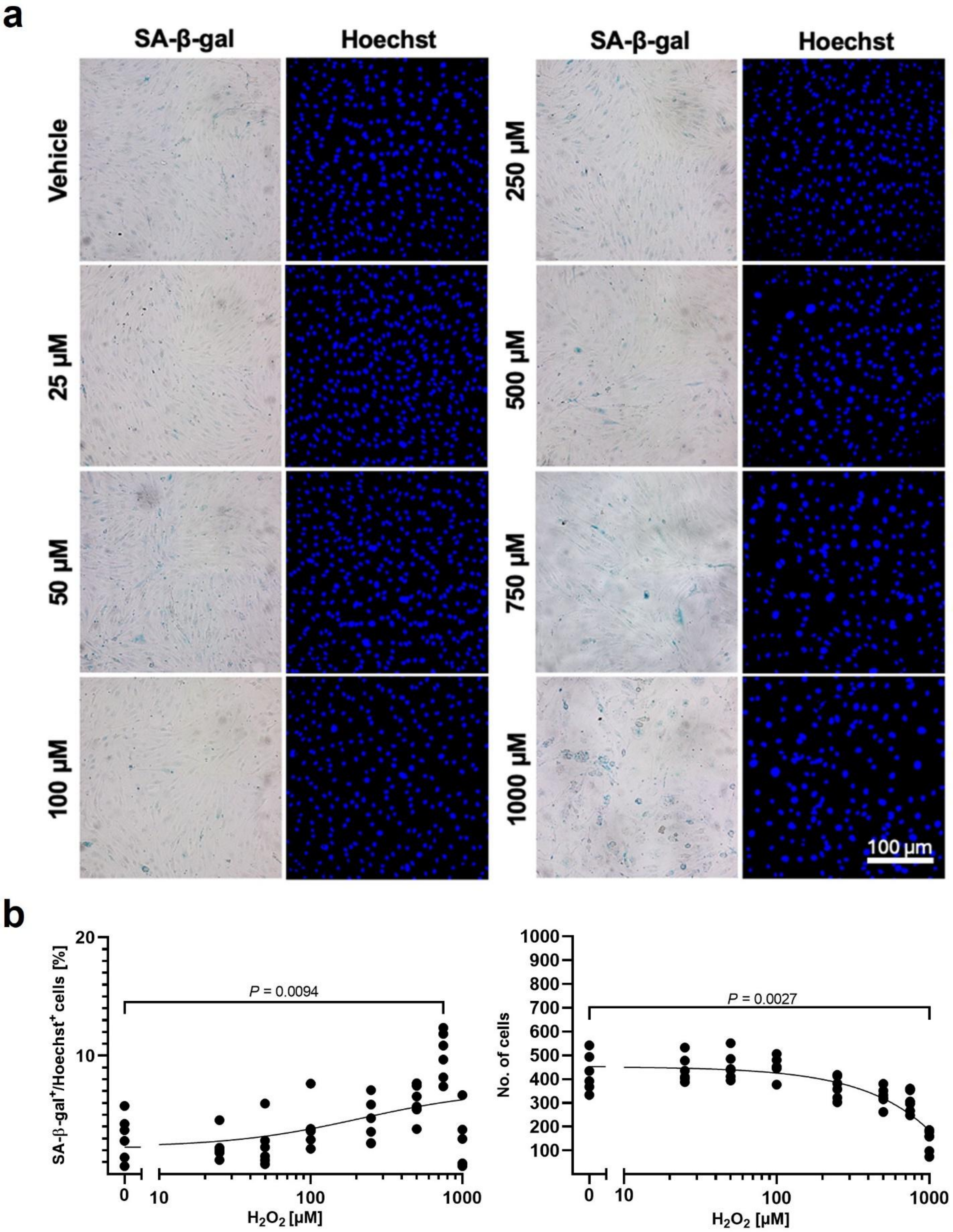


**Figure S3: Concentration-dependent increase in SA-β-gal activity after $H_2O_2$ treatment. A.** The SA-β-gal staining was imaged using bright-field microscopy. Fluorescence nuclear staining with Hoechst was performed to determine the total number of cells. **B.** The left graph shows the proportion of SA-β-gal positive cells at 72 hours of treatment. Kruskal-Wallis test: $\chi^2(8, N = 48) = 26.31$, $P = 0.0004$, $\eta^2 = 0.560$, followed by Dunn's post hoc test. The right graph represents the change in the total number of cells per concentration. Kruskal-Wallis test: $\chi^2(8, N = 48) = 34.42$, $P < 0.0001$, $\eta^2 = 0.732$, followed by Dunn's post hoc test.

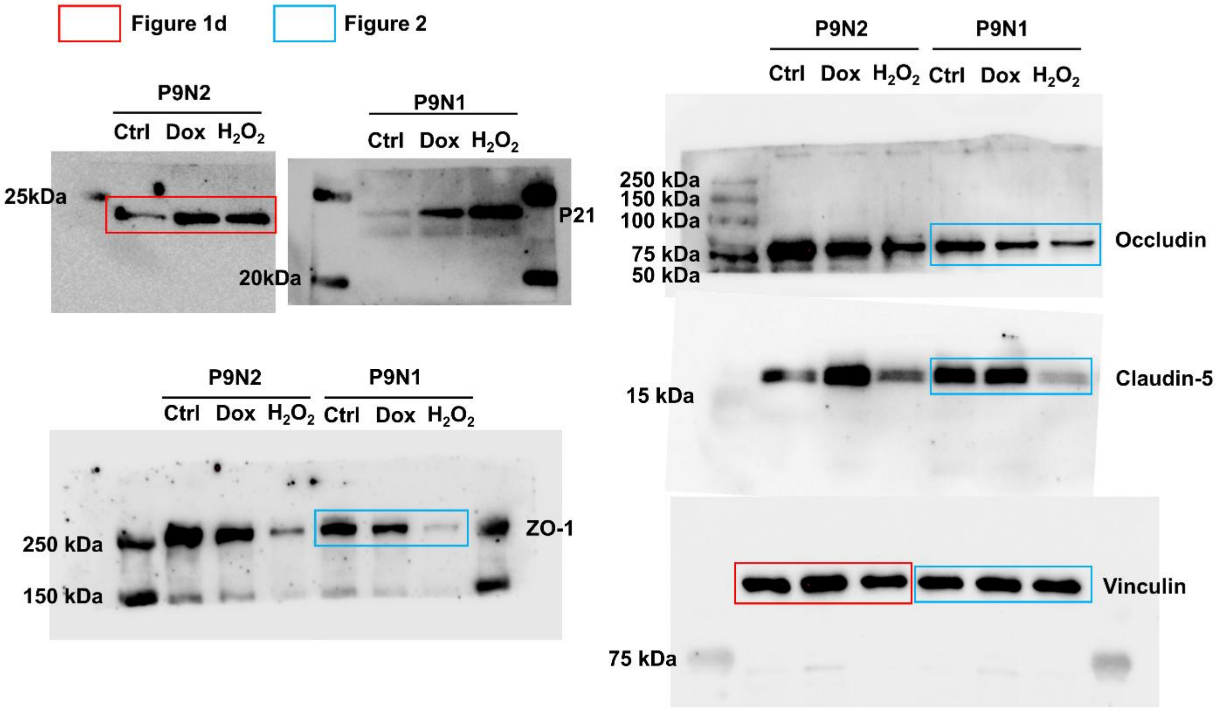


**Figure S4: Full western blots with size ladder.**

**Supplementary Table 1. List of primary antibodies used for immunoblot analysis.** IF = immunofluorescence, n/a = not available, WB = Western blot.

| Target | Clone, host species, type | Company (catalogue no.) | Dilution | RRID | Validation |
|---|---|---|---|---|---|
| Claudin 5 | 4C3C2, mouse, mAB | ThermoFisher (35-2500) | 1:1000 | AB_2533200 | WB: single band at ~18-23 kDa; validated in-house by IF (junctional localization) |
| Occludin | n/a, rabbit, pAB | Proteintech (13409-1-AP) | 1:8000 | AB_2156308 | Manufacturer knockdown/knockout validated; WB: ~59-65 kDa (multiple bands possible due to post-translational modification); validated in-house by IF (junctional localization) |
| P21 | 12D1, rabbit, mAB | Cell Signaling (2947) | 1:1000 | AB_823586 | WB: single band at ~21-23 kDa; widely validated in the literature |
| Vinculin | n/a, rabbit, pAB | Proteintech (26520-1-AP) | 1:20000 | AB_2868558 | WB: single band at ~116 kDa; standard loading control |
| ZO-1 | 1A12, mouse, mAB | ThermoFisher (33-9100) | 1:1000 | AB_2533147 | WB: single band at ~200-225 kDa; validated in-house by IF (junctional localization) |

**Supplementary Table 2. List of secondary antibodies used for immunoblot analysis.** HRP = horseradish peroxidase.

| Target (coupled to) | Company (catalogue no.) | Dilution | RRID |
|---|---|---|---|
| Mouse IgG (HRP) | ThermoFisher (A16017) | 1:10000 | AB_2534691 |
| Rabbit IgG (HRP) | ThermoFisher (A16035) | 1:10000 | AB_2534709 |
